# A Computational Framework to Study the Primary Lifecycle Metabolism of *Arabidopsis thaliana*

**DOI:** 10.1101/761189

**Authors:** Wheaton L. Schroeder, Rajib Saha

## Abstract

Stoichiometric Models of metabolism have proven valuable tools for increased understanding of metabolism and accuracy of synthetic biology interventions to achieve desirable phenotypes. Such models have been used in conjunction with optimization-based and have provided “snapshot” views of organism metabolism at specific stages of growth, generally at exponential growth. This approach has limitations in that metabolic history of the modeled system cannot be studied. The inability to study the complete metabolic history has limited stoichiometric metabolic modeling only to the static investigations of an inherently dynamic process. In this work, we have sought to address this limitation by introducing an optimization-based computational framework and applying to a stoichiometric model of the model plant *Arabidopsis thaliana* of four linked sub-models of leaf, root, seed, and stem tissues which models the core carbon metabolism through the lifecycle of arabidopsis (named as p-ath780). Uniquely, this framework and model considers diurnal metabolism, changes in tissue mass, carbohydrate storage, and loss of plant mass to senescence and seed dispersal. p-ath780 provide “snapshots” of core-carbon metabolism at one hour intervals of growth, in order to show the evolution of metabolism and whole-plant growth across the lifecycle of a single representative plant. Further, it can simulate important growth stages including seed germination, leaf development, flower production, and silique ripening. The computational framework has shown broad agreement with published experimental data in tissue mass yield, maintenance cost, senescence cost, and whole-plant growth checkpoints. Having focused on core-carbon metabolism, it serves as a scaffold for lifecycle models of other plant systems, to further increase the sophistication of *in silico* metabolic modeling, and to increase the range of hypotheses which can be investigated *in silico*. As an example, we have investigated the effect of alternate growth objectives on this plant over the lifecycle.

**Author Summary:** In an attempt to study the evolution of metabolism across the lifecycle of plants, in this work we have created an optimization-based framework for the *in silico* modeling of plant metabolism across the lifecycle of a model plant. We then applied this framework to four core-carbon tissue-level (namely, leaf, root, seed, and stem) stoichiometric models of the model plant species *Arabidopsis thaliana*, and further informed this framework with a wide array of published *in vivo* data to increase model and framework accuracy. Unique to the p-ath780 model, comparted to other models of plant metabolism, is the simultaneous considerations of diurnal metabolism, carbohydrate storage, changes in tissue mass (including losses), and changes in metabolism with respect to plant growth stage. This provides a more complete picture of plant metabolism and allows for a wider array of future studies of plant metabolism, particularly since we have only modeled the core carbon metabolism of *A. thaliana*, allowing this work to serve as a framework for studies of other plant systems.

## Introduction

The use of synthetic biology for the engineering of uni- and multi-cellular organisms to enhance desirable phenotypes in microbe, plant, and animal systems, has been well established and has been capable of affecting the lives of millions of individuals, such as in the case of artemisinin production in yeast or enhancing nutritional value of agricultural products [1–2]. Synthetic biology techniques have been applied to many plant systems such as tomatoes [3], rice [4], and maize [5] to produce enhanced phenotypes often with application to human nutrition [2], pest resistance [5], and resilience to abiotic stresses [6]. Many of these efforts have focused on a genetic understanding and manipulation of the plant system (or plant tissue) in question, having relied on intuitive interventions such as changes in regulation, insertion of new gene(s), and deletion of gene(s) from competing pathway(s) [2,5,6]. Alternatively, computation-based systems biology approaches, such as the use of stoichiometric genome-scale models (GSMs) of metabolism, have predicted non-intuitive genetic interventions [7] by accounting for Gene-Protein-Reaction (GPR) links and understanding how a gene knockout, or a change in gene regulation, affects the entire system through tools such as Flux Balance Analysis (FBA) [8], OptKnock [9], and OptForce [10]. Other tools are built upon previously existing tools, such as dynamic FBA (dFBA), which performs FBA over windows of time by solving a non-dynamic linear or a static linear problem, both of which integrate system variables over discrete time windows to solve to metabolite concentration, in addition to reaction flux [11]. Such tools have led to enhanced mechanistic understanding for exploring the system-wide effects of synthetic biology interventions especially in a microbial or a fungal system, such as *E. coli* [10], cyanobacteria [12], and yeast [13].

Stoichiometric global plant models, which treat the metabolism of the plant as a single unit, have been developed for *Arabidopsis thaliana* (hereafter arabidopsis) [14–17], *Zea mayz* (maize) [18], *Sorghum bicolor* (sorghum) [19], *Saccharum officinarum* (sugarcane) [19], *Brassica napus* (rapeseed) [19], and *Oryza sativa* (rice) [20]. These models have sought to analyze metabolic maintenance, response to abiotic stimuli, enzyme regulation changes, and metabolism as a whole at steady state (or pseudo-steady state). In addition, tissue-specific single-unit models have been reconstructed for various arabidopsis tissues [21], a maize leaf [22], and a barley seed [23] to better understand how present metabolites, metabolic pathways, and nutrient availability differ between tissues. Multi-tissue models have been created to characterize whole-plant metabolism for arabidopsis [16] and barley [17] and subsequently to study whole-plant metabolic response to the diurnal cycle and the source-to-sink relationship of leaves and seeds [16,17]. These studies either have considered metabolism at a single point [14,15,18–20], having taken a metabolic “snapshot” of a single point in growth time (often in the exponential growth phase) or have considered a single diurnal cycle [16]. This approach has been inherently limited in that metabolism is a dynamic and cumulative process. To clarify, metabolic state is dependent on both on external factors, such as availability of light, carbon sources, and availability of micronutrients, which these “snapshots” have captured, but also are dependent on metabolic history. These limitations have been inconsequential for single-cell systems in that laboratory apparatuses have held single-cell cultures at an exponential growth state; therefore, the “snapshot” approach has given good approximation of metabolism in these steady-state systems. In contrast, multi-cellular organisms, such as plants, will have passed through multiple and distinct stages of growth throughout its lifecycle [24], and the organism cannot be held at a steady state growth point. For this study, we have chosen *Arabidopsis thaliana* as the multi-cellular organism for several reasons. Firstly, since the advent of modern genetics, arabidopsis has served as a model plant species in that it has a small genome; therefore, arabidopsis has been well studied. Secondly, arabidopsis has a limited number of basic tissues which will have required the construction of a tissue-level model. Thirdly, arabidopsis has at least two distinct metabolic modes dependent on the availability of light. When studying the effects of a synthetic biology intervention on a plant system, such as arabidopsis, understanding the evolution of metabolism throughout the plant lifecycle can increase understanding of the cumulative effect of a synthetic biology intervention. The multi-tissue Edinburgh forest model, which has made use of Ordinary Differential Equations (ODEs) rather than stoichiometric matrices, has modeled the lifecycle of a tree for the purposes of studying lumber yield [16,17]; however, the intent of the aforementioned model has not been to consider individual reactions or genetic interventions, and therefore the GPR links which are central to the sought understanding and testing hypotheses when using SMs have not been included.

In this work, a core carbon stoichiometric metabolic model of arabidopsis has been reconstructed which consists of major primary carbon metabolism pathways, including, but not limited to, photosynthesis; the citrate cycle; starch and sucrose synthesis; fatty acid synthesis and degradation; and amino acid synthesis. The multi-tissue arabidopsis stoichiometric model, referred to as p-ath780 has 1033 total (and 633 unique) reactions (R), 1157 total (and 325 unique) metabolites (M), and accounts for 780 genes (G) including 42 chloroplastic and 11 mitochondrial genes. The model p-ath780 (**p**lant-scale primary **a**rabidopsis **th**aliana model including **780** genes) consists of four tissue-level models of metabolism: leaf (R: 537, M: 479, and G: 703), root (R: 130, M: 126, and G: 250), seed (R: 428, M: 411, and G: 529), and stem (R: 160, M: 140, and G: 250). The models are linked to one another and their respective environment by a comprehensive Flux Balance Analysis (FBA)-based [8] optimization framework [25] which considers both inter-tissue and environmental interactions. These four tissues have been chosen for model reconstruction to represent core plant functions. The root has been chosen and reconstructed for nutrient uptake and growth; the leaf for photosynthesis, carbon fixation, and as a source tissue for plant nutrition; the seed for metabolite storage and a sink tissue for metabolic investment; and the stem for metabolic transport and acting as a conduit for all metabolic interactions between other tissues. The dFBA method determines metabolite concentrations at the start and end points of the time frame [11], whereas the method developed does not focus on concentrations and considers multiple points within the time interval to make more accurate time-derivative estimates of steps in plant and tissue masses, as well as plant maintenance and senescence costs. The optimization framework of the p-ath780 model has taken a series of metabolic “snapshots” of arabidopsis metabolism throughout the lifecycle of a single representative plant subject to diurnal status, carbohydrate storage/uptake, changes in tissue mass (including losses), changes in relative tissues masses (due to growth stages), and changes in metabolism with respect to plant growth stage. p-ath780 has taken “snapshots” at hour intervals, and information from these snapshots have advanced plant and tissue masses forward one hour, when the next “snapshot” is taken. The series of “snapshots” produced by p-ath780 has given a framework for the investigation of the central metabolism of arabidopsis across its lifecycle. Several different objectives for this optimization-based framework have been investigated, with the default framework being the maximization of plant growth. Other alternative objectives investigated have included linear photonic efficiency and seed fatty acid production. This framework with the default objective has shown general agreement with experimental data and is potentially useful as an initial framework for other plant systems.

## Results

### *Reconstruction of arabidopsis* primary carbon metabolism in tissue-specific models

Figure 1 shows an overview of the workflow designed for determining the optimal reaction rates and mass step for each “snapshot” (top), how these “snapshots” have been advanced from one time point to the next; and how tissues have interacted at various stages of growth along with listing some important characteristics of a given growth stage that differs from other stages (bottom). In order to track the important metabolic interactions and transactions within and between major tissues of arabidopsis plant, namely seed, leaf, root, and stem, corresponding tissue-level metabolic models have been reconstructed. Model files for each tissue can be found in Supplemental Files 1 (seed), 2 (leaf), 3 (root), and 4 (stem). Figure 2 shows a summary of the distribution of model reactions across KEGG-defined pathways of each tissue model and an overview of reasons for reaction inclusion through confidence scoring (see Method section) [26]. Figure 2(A) summarizes the pathways common to all tissues, Figure 2(B) summarizes the pathways common to seed and leaf tissues, and Figure 2(C-G) graphically summarize the sources of reactions in each tissue model and p-ath780 as a whole through confidence scores (see methods section) [26]. First, the seed model has been reconstructed based on gene annotations and available MFA data [27] and then tissue model reactions have been distributed across five compartments based on literature evidence (see list of works cited in Supplemental File 5): extracellular space, cytosol, non-green plastid, inner mitochondria, and outer mitochondria. Next, transport and exchange reactions have been added to the model based on literature evidence (see list of works cited in Supplemental File 5) or modeling necessity to increase model connectivity [26]. The leaf model has been reconstructed using common reactions and pathways from the seed model and having added new pathways and functions essential to the major functions of the leaf tissue, such as photosynthesis [18]. In addition to the five compartments in the seed model, the leaf model contains chloroplast and thylakoid compartments. Similarly, by having extracted common reactions/pathways from the seed model, the root and stem models have been reconstructed. The root and stem models have been focused primarily on nutrient uptake (root) and transport (root and stem). Both these models contain necessary transport/exchange reactions to ensure model connectivity and to facilitate their roles in transport processes. The stem and root models have all the subcellular compartments present in the seed model. Once initial reconstructions have been accomplished, thermodynamically infeasible cycles in addition to atom and charge imbalances have been resolved [26] and tissue-specific biomass equations based on literature information have been defined [18,27,28].

**Figure 1.**
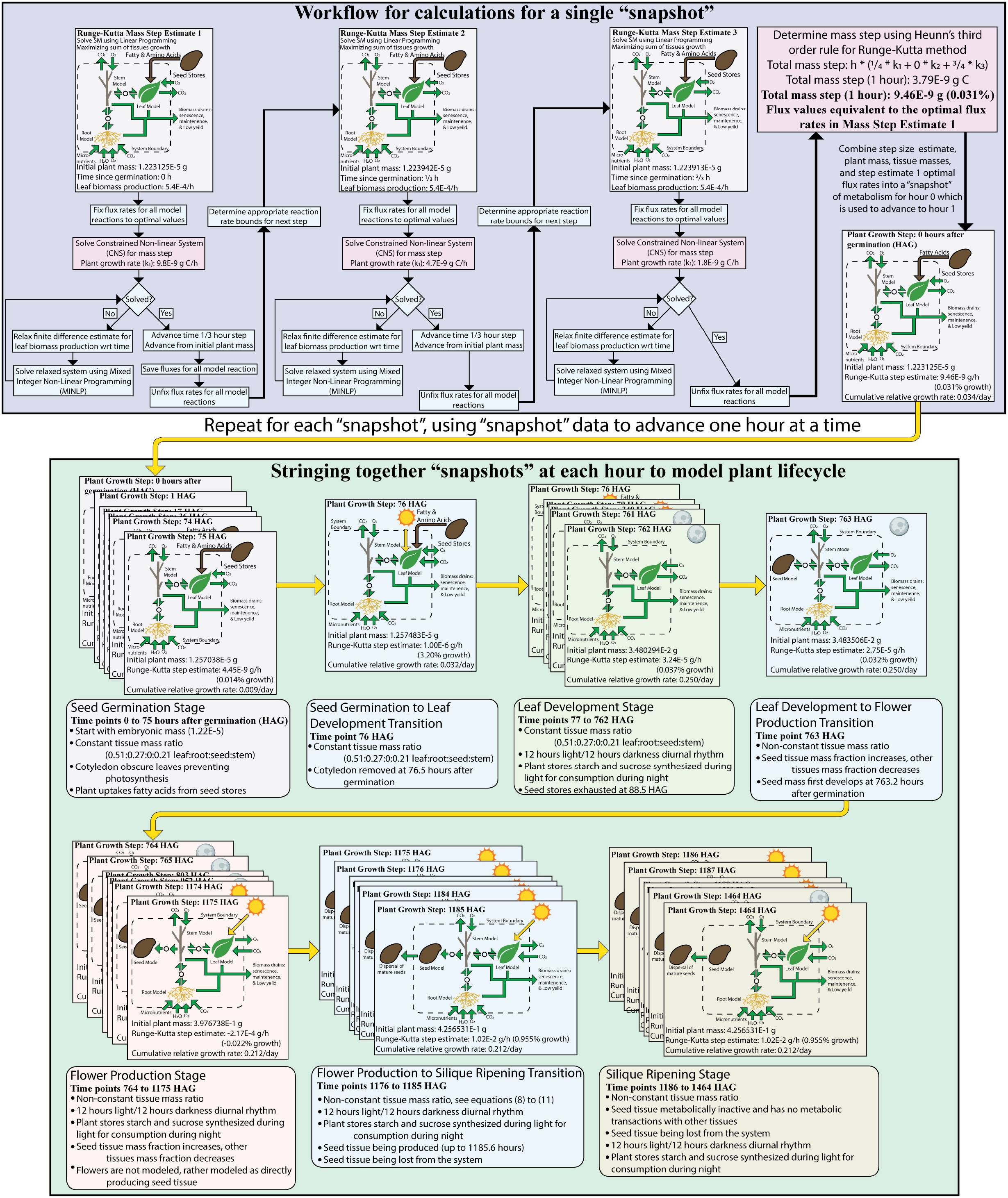
The design-build-test used cycle in constructing the p-ath780 model, where each box represents a step used in this cycle. The numbers in the lower left corner of each box indicates the approximate order in which these steps are undertaken for this cycle. This cycle is repeated until the *in silico* representation of *Arabidopsis thaliana* that is p-ath780 converges satisfactorily with *in vivo* experimental data as described in the results section.

**Figure 2.**
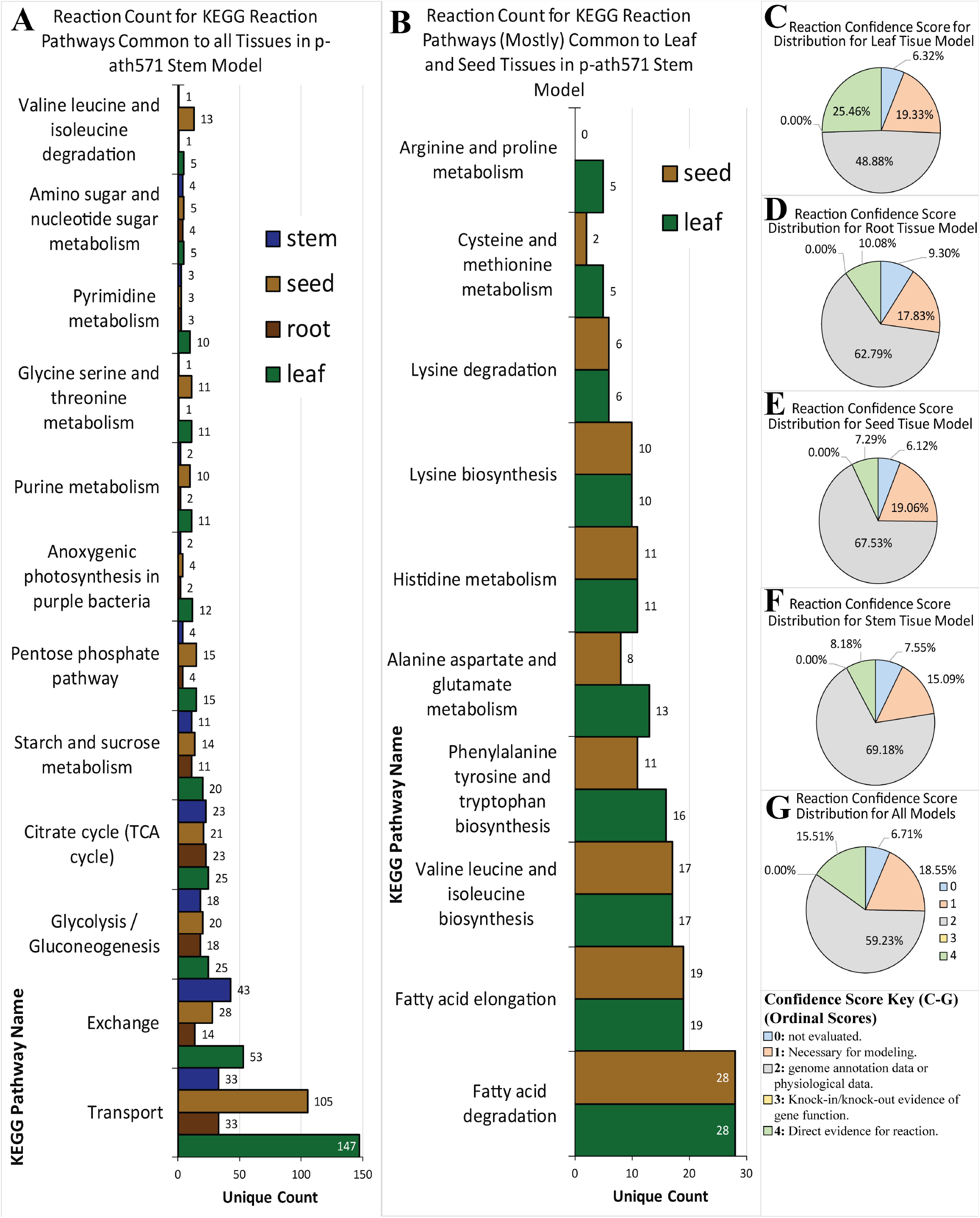
A heuristic look at the four tissue models in terms of number of reactions in various KEGG-defined pathways which provides some clarity as to the metabolic functions of each model (A and B) and in terms of the sources of included model reactions, indicated by confidence scores (C through F). A) A bar graph showing tissue model reaction counts in KEGG-defined pathways (with the exceptions being the user defined pathways of exchange and transport) common to most or all tissue (threshold: at least one model has at least three reactions in that pathway). B) Additional KEGG-defined pathways common to the seed and/or leaf model as these models contain more complete metabolism. C) – F) The source of each reaction included in the models through confidence scores for each tissue. G) The source of all reactions included in the p-ath780 model. See the methods section discussion related to confidence scores in this model.

### Development and tuning of the p-ath780 model

Once these core tissue models have been reconstructed and curated, these have been linked within a comprehensive FBA-based optimization framework (provided in Supplemental File 6) for *in silico* representation of metabolic behavior across the arabidopsis lifecycle. This framework has next been applied to the p-ath780 model that includes all four tissue-specific models and has 1033 total reactions, 1157 total metabolites, and 780 unique genes. Further details of the model development steps can be found in the methods section. The seed and leaf tissue have been selected to model an important source-to-sink relationship, whereas the stem and root tissues have been included to model nutrient transport and nutrient uptake in arabidopsis, respectively. The FBA-based framework has defined constraints related to tissue interactions and whole-plant growth heuristics based on experimental data, and also helped align *in silico* growth with experimentally determined *in vivo* growth through the modified design-build-test cycle shown in Figure 3, which will be discussed in greater detail later in this subsection.

**Figure 3.**
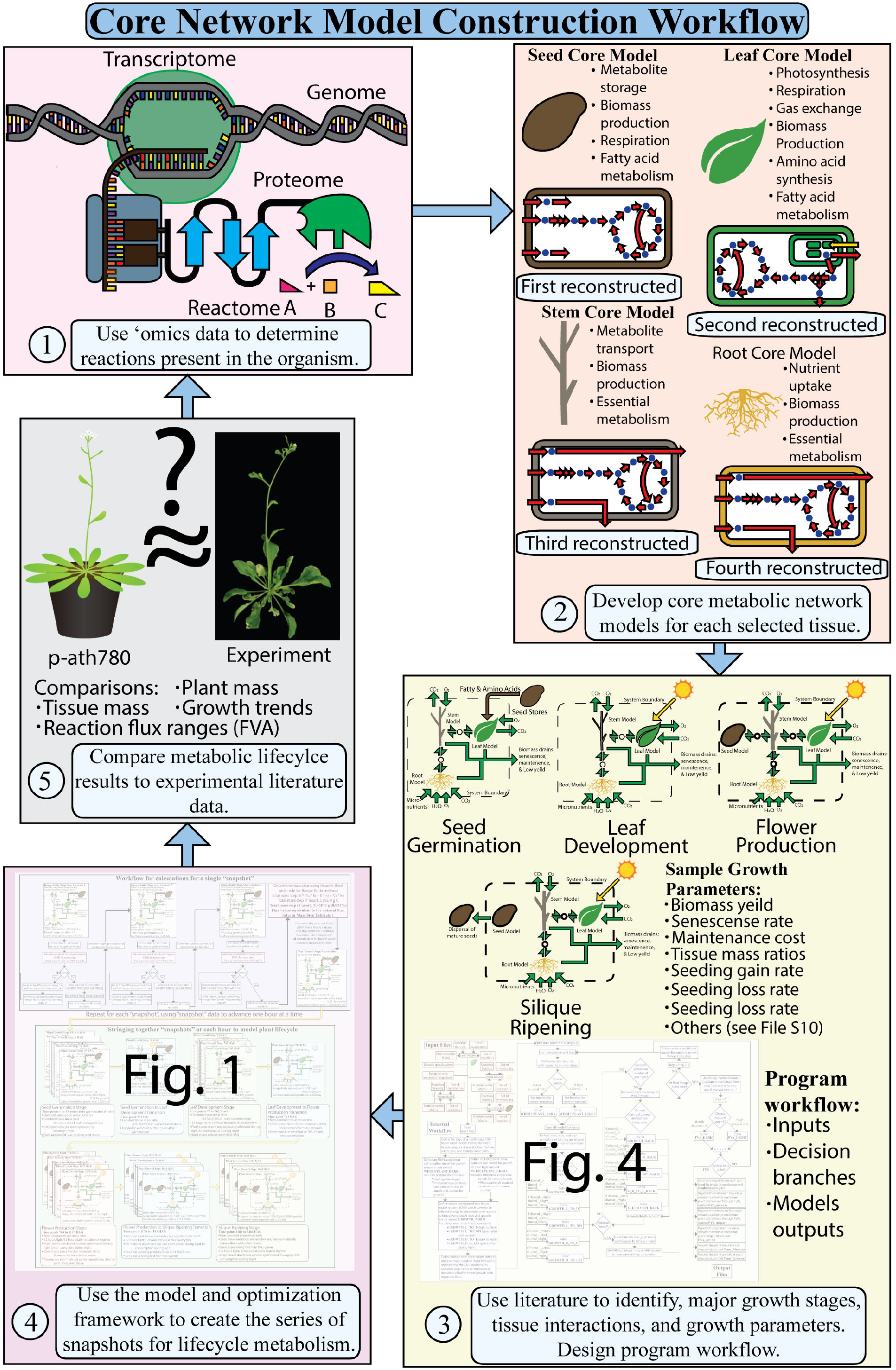
A simplified workflow of the calculations made to estimate the plant mass step size taken from one “snapshot” to the next (large top box) and visual representation of how these “snapshots” are strung together and grouped into stages and transitions (large bottom box). Each “snapshot” has been represented visually as small boxes containing initial time point, a figure highlighting major metabolic interactions, initial plant mass, step estimate, and cumulative relative growth rate. The contained figures show some major metabolic interactions across the plant system boundaries (full-headed arrows crossing the dashed system boundary) and indicates which tissues interact (single-headed arrows with circles to indicate a shared metabolic pool between the two tissues) for the given stage of growth which is indicated by the beveled box below the group of “snapshots”. The beveled boxes below a group of snapshots indicate the stage name (or transition name), the time points in growth which this stage encompasses, and some distinguishing characteristics of that stage.

The output of this framework has given metabolic “snapshots”, consisting of plant mass, growth rate, and flux rate of each reaction, at one-hour intervals across 61 days of growth, as the plant disperses all new seeds (through silique shattering) by 61 days after germination (DAG) [24]. After 61 DAG the plant begins to desiccate, eventually resulting in plant death [24]. The p-ath780 model is not used to model plant metabolism after 61 DAG because no *in vivo* data has been found in literature concerning the metabolomics of plant death and desiccation. This optimization-based framework has allowed for the sampling of changes in central carbon metabolism at different stages in the arabidopsis lifecycle (see Figure 3). For all the following analyses, the objective of this framework at each point has been the maximization of the sum of all tissue biomass production rates, unless otherwise indicated.

In having determined the mass steps taken for each hour intervals, three FBA-like calculations at 0, ⅓, and ⅔ hours past the hour have been made to increase the accuracy of the derivative estimate by an explicit numerical integration method calculating the mass step at each hour interval using Heunn’s rule for the third order Runge-Kutta method (see methods for greater detail). Figure 4 shows a more detailed workflow for each individual step in the form of a workflow diagram. Optimal flux points at every whole hour have been saved as the optimal flux rates at that growth point. To evaluate these balanced flux estimates, Flux Variability Analysis (FVA) [29] has been performed, at nine points, selected to represent each non-transition growth stage and diurnal status in those stages, subject to all growth constraints and a growth rate equivalent to the optimal growth rate (see methods for enumeration of these points).

**Figure 4.**
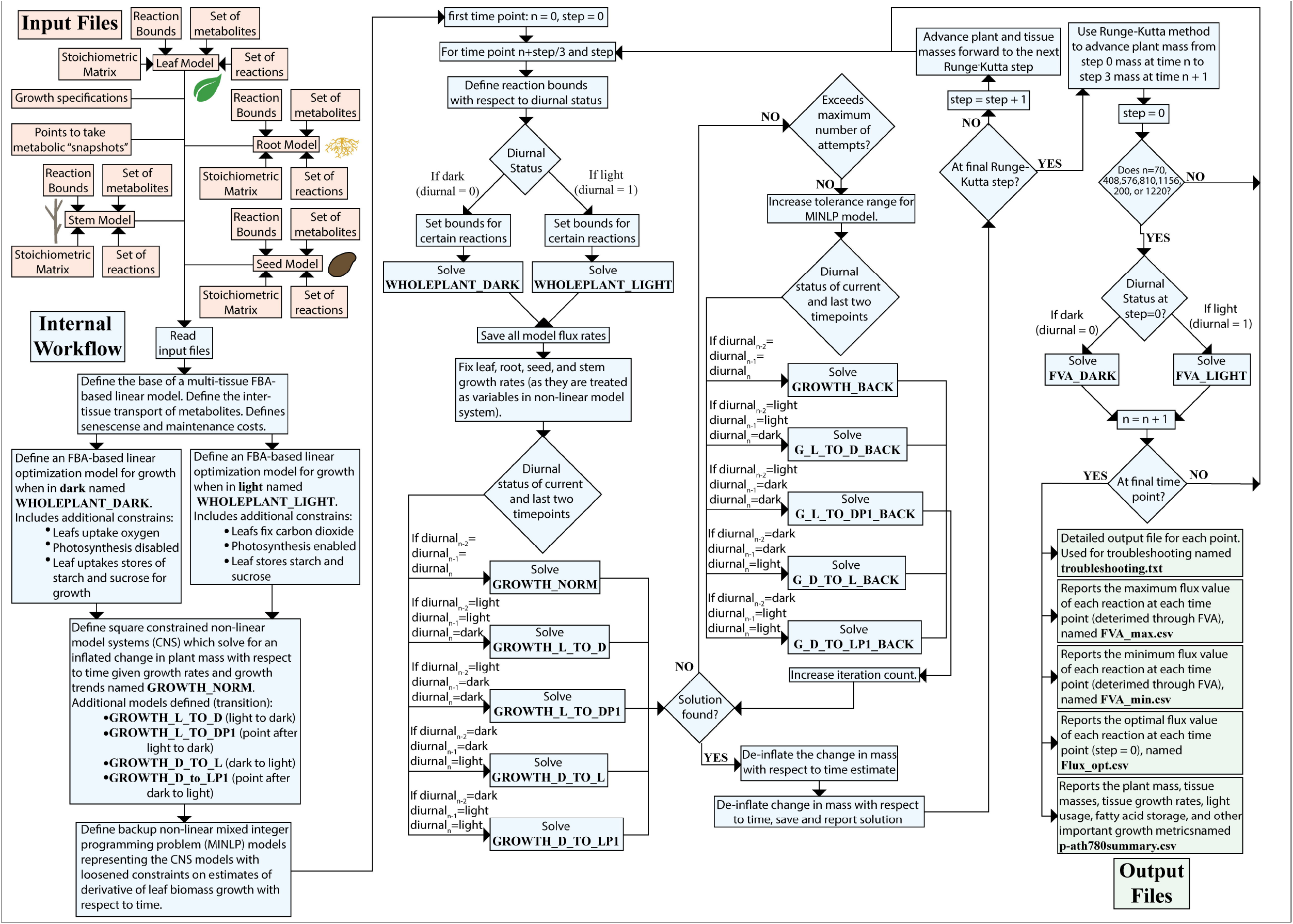
The workflow diagram of the p-ath780 model, including inputs (orange), outputs (green), and internal workflow (blue). The inputs for the p-ath780 model include each individual tissue model, a file of growth specifications, and a list of point at which to take metabolic “snapshots”. The internal workflow has read these inputs and then used them to construct model objects (bold, Times New Roman text) which are used to perform FBA, to solve for what the plant mass step is from the current to the next “snapshot”, and to perform FVA. For each iteration, the time of the snapshot is stepped forward 1/3 step (hour), the FBA model object is solved, the mass step is calculated, and process is repeated. Every third iteration (e.g. where step = 3), Heun’s method for a third order Runge-Kutta is used to estimate the plant mass step from the previous whole hour to the next whole hour and FVA is performed on the model at the previous whole hour using saved values. Once all iterations are complete (e.g. model is at final time point), then the output files are written.

The simulations of the p-ath780 model has been advanced through several growth stages using time point for changes in growth stage taken from experimental data [24]. In the seed germination stage, uptake of fatty and amino acids from seed storage has been modeled as a constant rate of fatty acid usage which results in all stored fatty and amino acids being depleted by the end of the seed germination stage [30]. The 12 hours of light and 12 hours of dark diurnal rhythm has been chosen to match experimental conditions for studies on starch and sucrose storage/uptake dependence on the diurnal cycle [31]. These patterns have been fit to a sine wave model constraint with ±1% tolerance. In growth stages when plant tissue ratios have been constant, the tissue mass ratio values had been taken from values typical for herbaceous plants [32].

Returning to the design-build-test cycle used to improve the p-ath780 model, experimental data related to plant growth and plant growth stages have been collected from a variety of literature sources to serve as checks for the accuracy of the modeled system [24,33]. The first set of experimental data has included mass data, including whole plant and individual tissue. At approximately 17, 24, and 31 DAG the total dry plant mass should be between 0.5 and 2.0 mg; 2 and 5 mg; and 10 and 30 mg, respectively [33]. Once the design-build-test cycle has been completed, the p-ath780 model has shown a total dry plant mass of 0.554 mg at 17 days (408 hours), 3.74 mg at 24 days (576 hours), and 25.2 mg at 31 days (744 hours) after germination, demonstrating growth consistent with *in vivo* data. Furthermore, the relative growth rate for the first 31 days of plant growth has been reported as between 0.21 and 0.25 day^−1^ [33], and the final p-ath780 has shown a relative growth rate of 0.246 over this time period. To adjust model behavior in latter stages of growth, tissue-specific mass data has been obtained from literature. Specifically, the dry weight of the stem, the leaves, and the seeds has been reported as approximately 188 mg (standard deviation 39.3 mg), 163.7 mg (standard deviation 52.0 mg), and 127.9 mg (standard deviation 52.7 mg), respectively [24]. As p-ath780 models both plant growth and loss of seed (and other) mass in the silique ripening stage, the peak mass of each of these tissues has been comparted to this data. In the final p-ath780 model, the peak mass of the stem, leaves, and seeds has been determined as 189 mg, 177 mg, and 130, respectively, all of which are within one standard deviation of the experimental value (see the methods section for how tissue masses are determined). In summary, through the results of the design-build-test cycle implemented, *in silico* tissue and plant mass values are similar to *in vivo* data, thus showing strong agreement with respect to growth trends.

In early rounds of model reconstruction, it has been noticed that the plant model’s photosynthesis is too efficient at fixing carbon. This is due to the fact that plants do not make full use of available light source(s), but the reconstructed metabolic model had been. Published *in vivo* data which has been used in the modeling and verification of p-ath780 made use of fluorescent lights, which have tight transmission spectra peaks at 544 and 609 nm [34]. In contrast, peak absorbance for plant leaves is at approximately 440 and 680 nm [35]. The problem of the availability of light has been addressed by scaling the transmission of the fluorescent lights by the absorbance of plant leaves. This has left approximately 21.06% of light transmitted by the fluorescent bulbs usable by the plant (see Supplemental File 5 and methods). An additional restriction, namely biomass yield, has also been placed on metabolic efficiency in plant systems. Biomass yield has been defined as the carbon fraction of biomass produced appearing in new growth for each unit of carbon used for growth [36]. This yield value accounts for repair of existing biomass and replacement of lost biomass. Experimentally, this value has been identified as generally between 0.7 and 0.85 [37]. Here, for p-ath780 mass values to align with experimental data, two separate mass yield vales have been set at 0.32 and 0.23 for when the plant system lacks and has seed tissue respectively. This represents an incongruity with experimental evidence, although this value is still in the same order of magnitude as experimental evidence. All files necessary for p-ath780 have been included with this work in Supplemental Files 7 through 16. The *in silico* results of the final p-ath780 model can be found in Supplemental File 17.

### In silico *Plant Growth under* Alternative Objective Functions

A total of six different objective functions for p-ath780 have been investigated and a summary of that investigation has been shown in Table 1. In all cases, root and stem tissue objectives have been defined as biomass production, where the leaf and seed objective functions are varied. When any tissue has a non-biomass objective, that objective is weighted by some scaling factor (either α for light-based objectives or β for fatty acid-based objectives) to ensure the new terms do not dominate or be insignificant compared to biomass (e.g. be an order or magnitude or more different) and to investigate the different effects of weight factors. The first row (green) of Table 1 contains the *in vivo* arabidopsis data which has been used as targets and verification of the p-ath780 model. The second row (blue) contains the *in silico* data from p-ath780 where the objective for all tissues is biomass production (default objective), and has summarized the findings of the preceding subsection. For the mathematical definition of this and other objective functions discussed here see the methods section. The next two objective functions presented (grey) we have considered set the seed tissue-level objective as the maximization of fatty acid stored in the seed tissue at two different weight factor values (β). At low β values, this causes a scavenging of carbon wasted in the plant metabolism which is then diverted to the seed fatty acid production without a change in plant growth. At high β values, this alternate seed objective results in stunted plant growth as carbon used elsewhere is diverted to the seed tissue. This alternate objective function has no effect when the plant does not have seed tissue present. Photonic efficiency for the leaf tissue has been attempted as an alternative objective function and results are reported in the next two grey rows; however, depending on the weight of the photonic efficiency parameter, the model is generally photophobic (no light has been uptaken) or grows as normal. In all photophobic growth cases, the plant mass of the model eventually becomes negative, leading to nonsense in later time points, thus the results of these α values are not reported. No α value has been identified which produced a result between these extremes (photophobic and normal growth), and these attempts are not included in this work. The full output of each reported result can be found in Supplemental File 17. The final objective function investigated (and reported in Table 1) is one which combined the leaf linear photonic efficiency objective and the seed fatty acid storage objective at a moderate weight value (values enumerated in Table 1). As with other linear photonic efficiency objectives, the amount of light uptaken by the plant is unaffected, and as with other investigations of the seed fatty acid objective, the plant growth is stunted when the seed tissue is present. In summary, the p-ath780 model is robust to small and moderate perturbations in the objective related to photonic efficiency, fails with large perturbations to photonic efficiency objectives, and results in continuously changeable growth levels to metabolite production objectives.

**Table 1:**
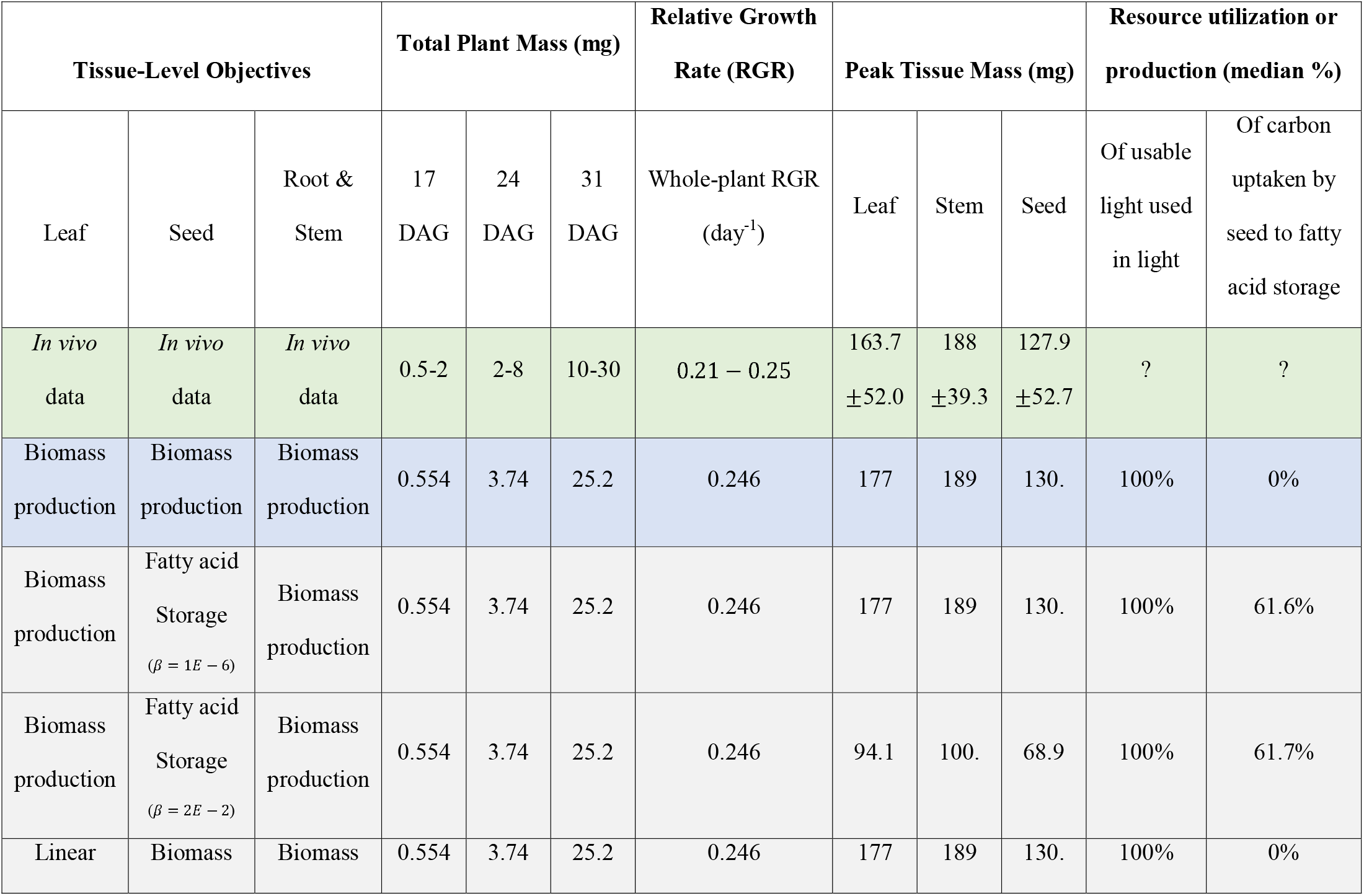
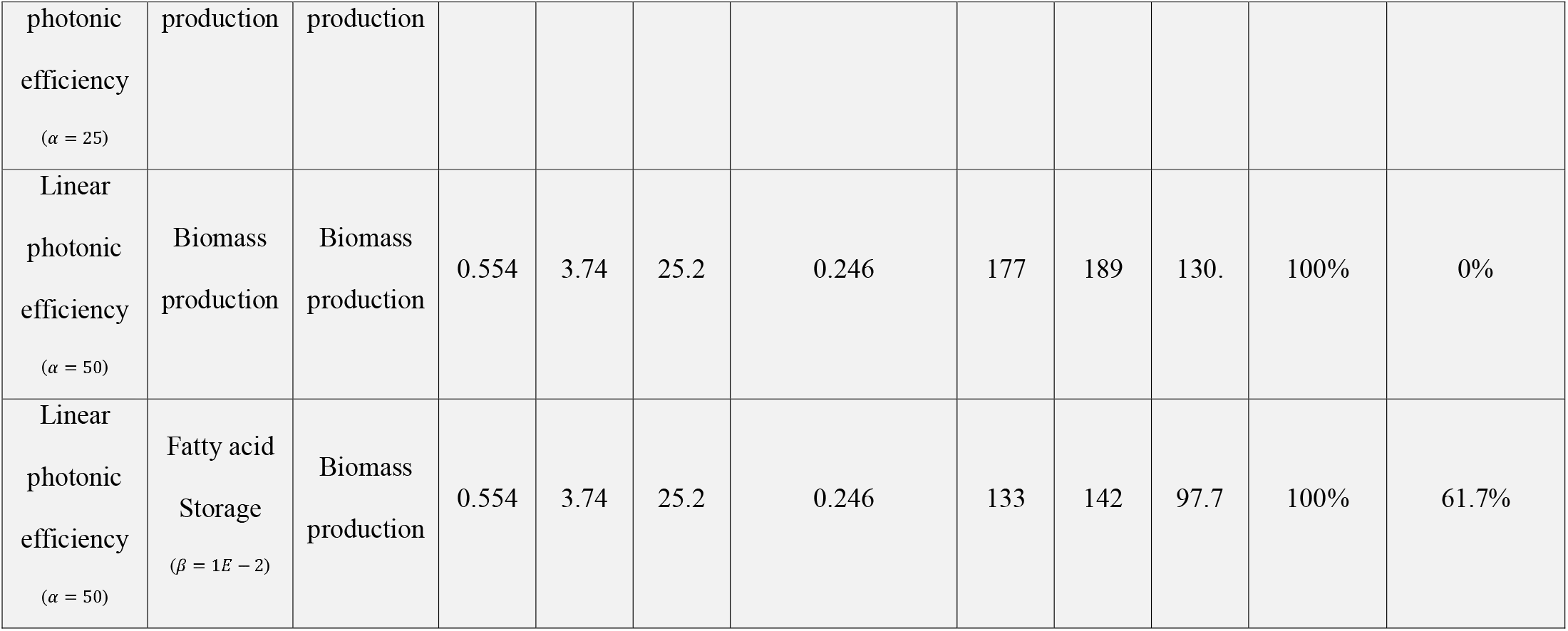
This table compares the critical mass-based metrics which have been used for *in silico* to *in vivo* data comparison across different objective functions for the p-ath780 model. The final two columns are for metrics which have been targets of change when applying different objective functions. The green row is the *in vivo* data ranges which have served as targets for *in silico* model behavior. The blue row is the behavior of the p-ath780 model with the usual objective function used in all other analyses (maximizing biomass production of all tissues). The grey rows are alternate objective functions we have explored. The photonic efficiency objective functions for the leaf tissue have caused lower fractions of available light used, while the fatty acid production objective for the seed tissue has caused greater diversion of carbon resources toward fatty acid production and storage. All alternative objective functions have resulted in lower mass production *in silico*.

## Discussion

In the current work, a multi-tissue core metabolism stoichiometric model, including leaf, root, seed, and stem tissues, of *Arabidopsis thaliana* has been reconstructed (Figure 3), and linked in an FBA-based optimization framework (Figure 1). This framework has been embedded in a workflow (Figure 1) which has simulated how plant metabolism evolves over time with respect to the presence or absence of light, the transition to different growth stages, and the gain or loss of tissues (such as seed). This model has incorporated a wide variety of data which has not been incorporated in other stoichiometric modeling efforts such as the effect of plant mass, the effect of tissue mass difference on tissue interactions, whole-plant growth heuristics such as yield, the availability of usable light, and biomass-based plant maintenance (as opposed to ATP-based). The tissue models taken together with these literature-based constraints has been named the p-ath780 model. The whole-plant growth characteristics of p-ath780 have shown general agreement with experimental data, particularly with respect to whole plant mass at certain growth milestones and lifecycle tissue yields.

The design-build-test cycle used to develop and tune p-ath780, shown in Figure 3, has been implemented. As a result, in the final p-ath780 model, *in silico* predictions compared well to *in vivo* data, particularly plant, leaf, seed, and stem masses, with the exception of biomass yield. The incongruity between *in* vivo and *in silico* biomass yield has likely resulted from the p-ath780 model only having included primary carbon metabolism, which in turn means that plant biomass has been built entirely from generally less metabolically expensive primary metabolites. This had resulted in too efficient biomass production, hence the lower yield for the model. This discrepancy in biomass yield has served to highlight the large effect of secondary metabolism on plant growth and has served as a correction factor on the model due to the lack of modeled secondary metabolism. In addition, likely the plant mass yield is lower when seed tissue is present because flower tissue is metabolically expensive yet is not modeled in this work. In additional, biomass drains for plant senescence and maintenance have been included [36,37].

Once the model has been developed, six different objective functions have been applied to it, including the default objective of maximizing plant growth, linear photonic efficiency, and seed fatty acid production. In summary, the p-ath780 model is robust to small and moderate perturbations in the objective related to photonic efficiency, breaks when large perturbations are made to the photonic efficiency objective, and is capable of some fine tuning with respect to metabolite production objectives. The behavior of p-ath780 with respect to the linear photonic efficiency objective function is due to multiple factors. First, as the partially photophobic case exists, this suggests that seed tissue is the most metabolically expensive tissue to create. This is as expected because the seed tissue requires storage of high-energy molecules such as fatty acids, proteins, and sugars to feed its embryo when dispersed. Further, the rate of biomass production for all tissues are linked in the optimization-based framework. Secondly, seed tissue has a target fraction of overall plant mass which it must grow to for each hour interval of the flower development stage. If seed tissue is too metabolically expensive to produce, relative to the cost to uptake more light, it appears that the solution strategy then becomes to decrease the mass of other tissues while leaving the growth of the seed tissue to be minimal. This can result in sharp changes in mass which falls outside the realm of stability for Heunn’s third order Runge-Kutta rule, resulting in predictions with no biological relevance. Thirdly, biomass composition and metabolic cost is not dependent on the amount of light uptaken, so the biomass cost is constant with respect to light uptaken so there is no steady equilibrium between the two terms except at the extremes. Fourthly, minimizing light uptake and maximizing biomass growth as objectives are competitive, increase light uptake results in increased growth in stoichiometric models. In contrast, this is not an issue for maximizing fatty acid production as biomass partially consists of fatty acids, therefore these two objectives can be complimentary to some degree and a variety of β values can be used without the model failing to find a solution. Theoretically, there exists some highly-specific value of α at which the cost to the light needed to drive growth is balanced with the rate of production of new biomass, but this is an unsteady equilibrium which when the value of α is slightly perturbed finds the new equilibrium at either extreme. Therefore, the value of α might be imagined as a fulcrum between the two terms as illustrated in Figure 5. Figure 5(A) restates the linear photonic efficiency objective equation, and Figure 5(A-D) represents the action of α as a fulcrum. Figure 5(D) in particular illustrates why the p-ath780 model is robust to changes in the value of α below this theoretical balance point.

**Figure 5.**
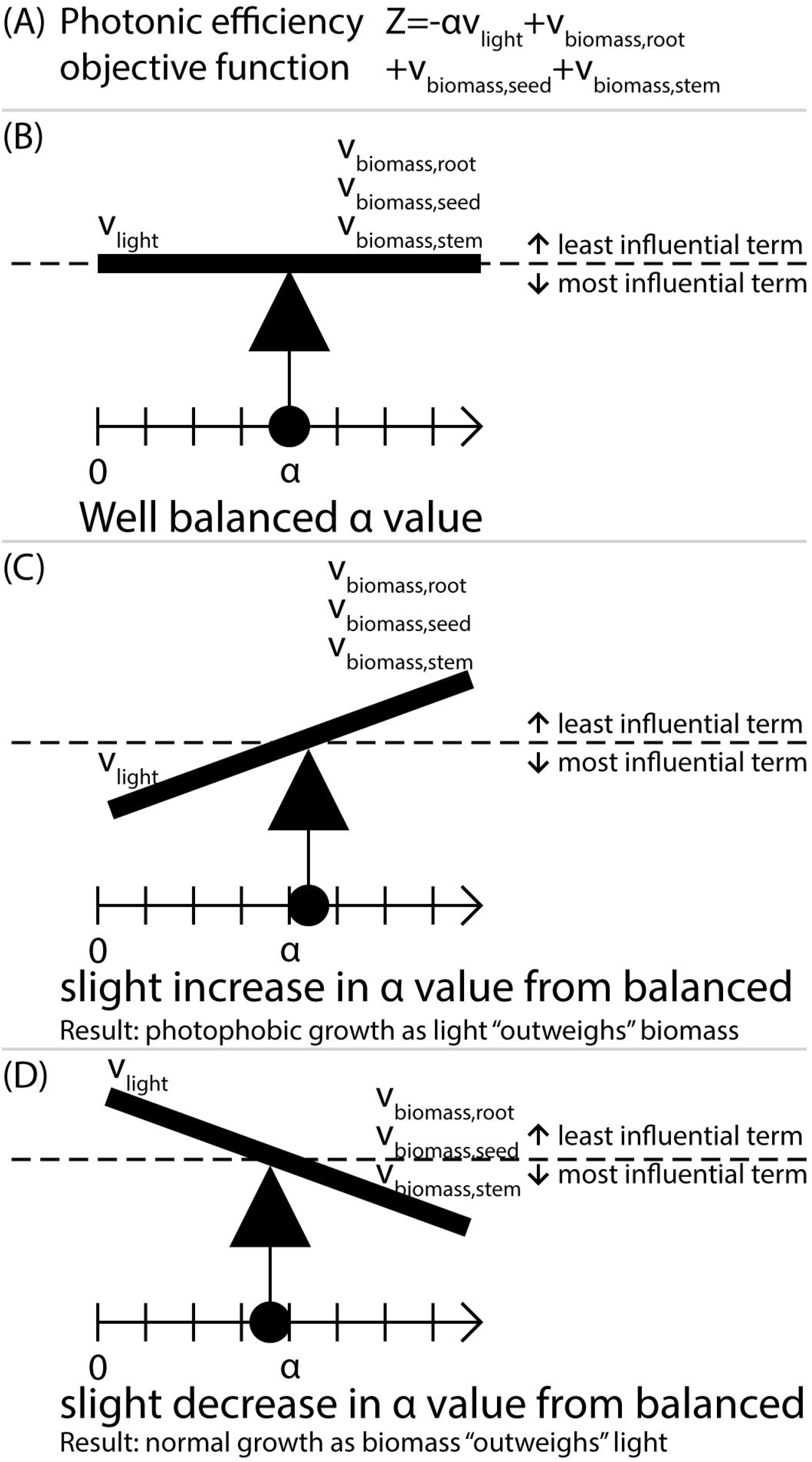
This figure highlights how the weight factor, α, fails to reach some kind of equilibrium between light uptake and biomass production in the linear photonic efficiency objective. A) Restates the linear photonic efficiency objective function. B) Shows the theoretical balance which might exist between light uptake and biomass production at some highly specific value of α. C) Shows how as slight increase in α from that point causes light to outweigh biomass in terms of influence on the objective function value. This results in photophobic growth as light “outweighs” growth. D) Shows how as slight decrease in α from that point causes light to outweigh biomass in terms of influence on the objective function value. This results in normal growth as growth “outweighs” light.

This work does not account for diurnal rhythms in the transcriptome of *Arabidopsis thaliana* for several reasons. Firstly, the majority of transcriptomic studies have focused on the regulatory network of proteins which regulate metabolism based on the availability of light and rhythm [43,44], rather than considering metabolic proteins which are represented in the p-ath780 model. Secondly, tissue-specific diurnal transcriptomic information is only available for the leaf tissue [43,44]. Further, these experiments generally consider a single point in the growth cycle of arabidopsis under specific growth conditions. The framework of p-ath780 is already highly constrained, and that the inclusion of too much data will invariably cause model failure. This is because *in vivo* experiments, in general and those used in this work, often occur under different conditions, at different points in the plant lifecycle, have different methods to some degree, or even seem quantitatively difference due to the noise inherent to biological systems, making alignment of quantitative *in vivo* data from too many sources impossible. In this work, we have decided to use data which described a wide range of time point in arabidopsis growth, such as biomass yield, relative growth rate, growth up to a certain time point, and overall tissue yield, rather than data which may be specific to a single point in growth, such as transcriptomics.

This work provides the basis for much future development and sophistication. For instance, the current p-ath780 model could be further sophisticated by adding the secondary metabolism of the plant system, which is a considerable resource drain in many plant systems. Further, at present several simplifications are made regarding tissues, particularly related to seed tissue. For instance, the model currently assumes that when the plant is flowering, that flower biomass and metabolism is roughly equivalent to that of the seed. While this resulted in a simpler model, this model cannot be then used to investigate certain metabolic hypothesis such as the cost to the plant resulting from flower pigmentation, pollen, and nectar production. Future work will include producing models for other plant tissues, such as flowers. In addition, as this is a core carbon metabolism model, it is likely quite similar to the core metabolism of other plant systems; therefore, the p-ath780 model can serve as a basis for the development of lifecycle models for other plant systems, particularly annual eudicots which are of agricultural interest, such as rice (*Oryza sativa*), potatoes (*Solanum tuberosum*), tomatoes (*Solanum lycopersicum*), and soybeans (*Glycine max*).

## Methods

### Overview of the reconstruction of core metabolic models of leaf, root, seed, and stem tissues. The seed tissue model

The general workflow which has been used for the development of the four core tissue models has been illustrated in Figure 3. We have developed the seed model first, with the central metabolic pathways based on a Metabolic Flux Analysis (MFA) of four seed genotypes published previously [27]. We have then manually filled gaps in this model with reactions based on literature and genomic evidence [26] or with reactions being necessary for ensuring model connectivity. The stoichiometric coefficients of biomass precursors have been determined using sink reactions, dry biomass weight composition, and amino acid mass ratios provided in a previous work [27] (see Supplemental File 22). The resultant seed tissue model has focused on storage, respiration, and growth, and consists of 428 reactions, 529 genes, and 411 metabolites (included as Supplemental File 1).

### The leaf tissue model

Next, we have reconstructed the leaf model by taking common reactions/pathways from the seed model and adding synthesis pathways for amino acids that are not synthesized in the seed, in addition to photosynthesis, carbon fixation, gluconeogenesis, and transport reactions. We have then developed the biomass equation for the leaf tissue using that of a previously published Arabidopsis model [18] (see Supplemental File 22). The resultant leaf tissue model has focused on photosynthesis, respiration, gas exchange, fatty acid synthesis, and growth, and contains of 537 reactions, 703 genes, and 479 metabolites. We have included the leaf model with this work as Supplemental File 2.

### The root and stem tissue models

We have constructed the root and stem models, similarly, by extracting common reactions/pathways from the seed model and adding necessary transport and exchange reactions. Then exchange reactions have been added to allow the root to be linked to micronutrient uptake processes from the soil, and the stem to be involved in inter-tissue transport processes. In the absence of Arabidopsis-specific estimates, the dry weight composition of switchgrass (*Panicum virgatum*) root and stem [28] have been assumed to be equivalent to the biomass composition of these tissues in Arabidopsis. Due to the low detail level of the dry weight composition analysis, the biomass of root and stem tissues have been composed entirely of carbohydrates. The resultant root tissue model has focused on nutrient uptake, transport, and growth, consisting of 130 reactions, 250 genes, and 126 metabolites, while the stem tissue model focuses on transport and growth, consisting of 160 reactions, 250 genes, and 140 metabolites. We have included the root and stem models with this work as Supplemental Files 3 and 4 respectively.

### Confidence scoring

Reaction confidence scores have been defined in a manner consistent with a previously published protocol [26]. Additional information on confidence scoring of the p-ath780 model can be found in Supplemental File 22.

### Curation of all these tissue models

For all four models, we have balanced (both in terms of elements and charge) all model reactions and have resolved thermodynamically infeasible cycles by removing reactions, breaking composite reactions, and adding metabolic costs to transport reactions. For all these tissue models, GPR links have been established through a largely automated workflow utilizing the KEGG API for the majority of reactions using the code included in Supplemental File 18. This is followed by having manually curated the GPR links and/or inclusion rational of reactions with non-KEGG identifiers. The count of tissue model reactions present in KEGG-defined pathways is shown in Figure 2(A), showing pathways common to most all tissue models, and Figure 2(B), showing pathways common to seed and leaf tissues, these figures have been created using code included in Supplemental Files 19, 20, and 21. The results of this automated workflow can be found in Supplemental File 5. Sources for reactions included in leaf, root, seed, and stem models are shown in Figure 2(C-F), respectively through confidence scoring (see confidence score section). Similarly, the confidence scores for all reactions in the p-ath780 model have been reported in Figure 2(G).

### Overview of the developed of the optimization-based framework of p-ath780

The models have next been linked using well-known computational framework known for modeling microbial communities [25]. An objective function for each of these models has then been defined, specifying the maximization of the tissue-level biomass production rate followed by adding constraints for simulating growth in light and dark conditions. Next, literature information including embryo mass [30], initial tissue masses [38], growth stages [24], time points at which growth stages occur [24], constraints to link tissue growth rates to appropriate tissue ratios, transpiration [33,39], leaf surface area [28], usability of provided light [31,34,35], and defining changes in tissue mass ratios [24,40] has been integrated into these models, which are typically overlooked in most other SMs. In this work, we have decided to simulate arabidopsis biomass across 61 days (1464 hours) of growth, as all plant seeds are dispersed by day 61, and after which *in vivo* data on plant growth and mass is sparse. More specific details can be found in the following sub-sections. The full optimization-based framework used in this work has been provided in Supplemental File 6, and further requires Supplemental Files 7 through 16.

### Generalized statement of the FBA-like optimization-based framework used

The optimization-based framework used can be stated in general terms as follows. The FBA-based framework which determines the optimal rates of flux through each reactions can be stated as follows (using the default objective).

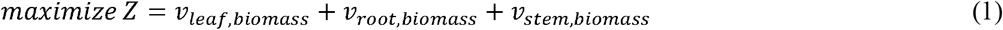

Subject to

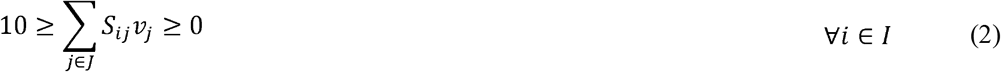

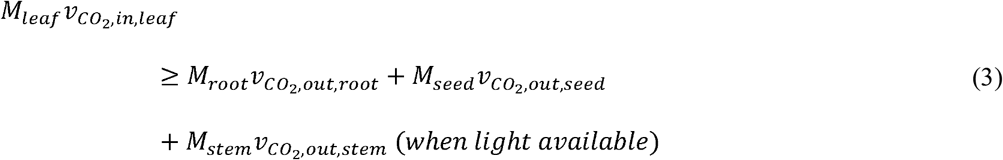

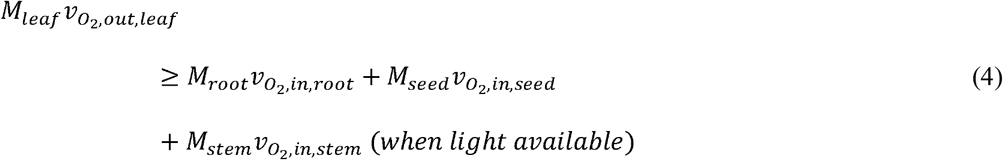

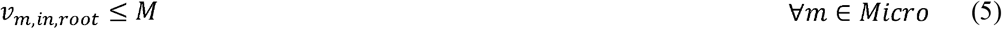

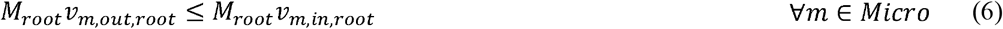

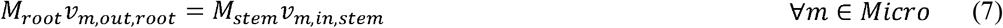

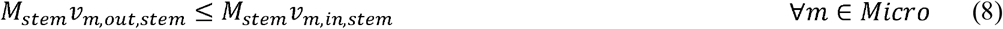

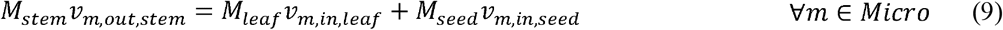

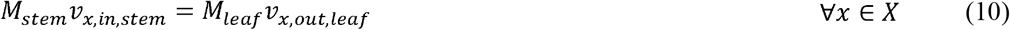

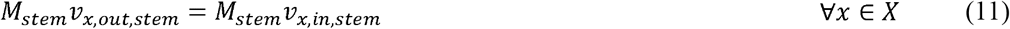

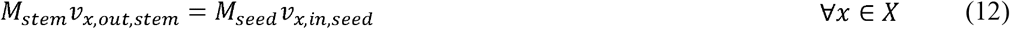

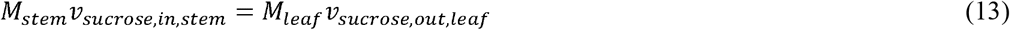

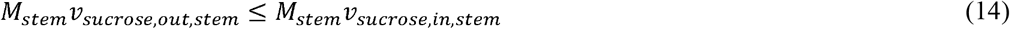

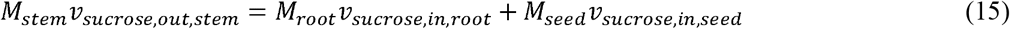

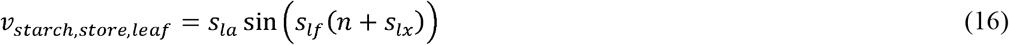

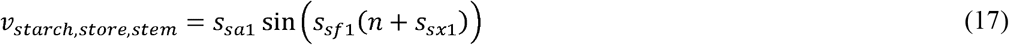

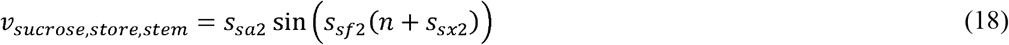

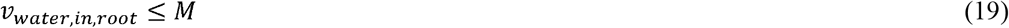

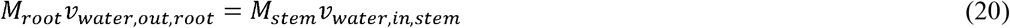

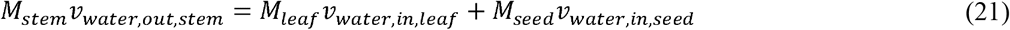

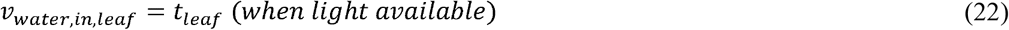

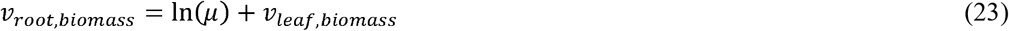

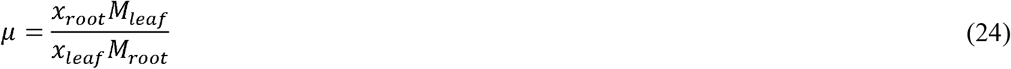

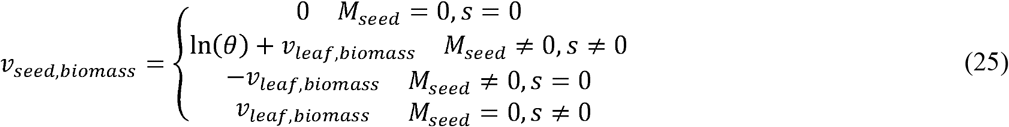

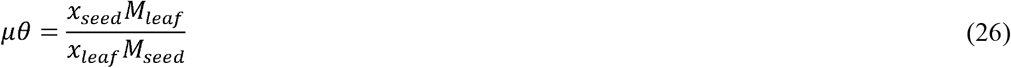

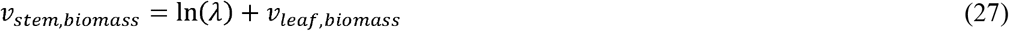

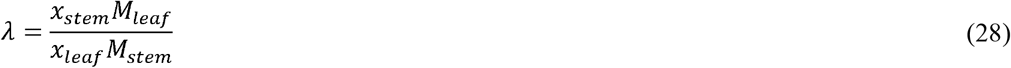

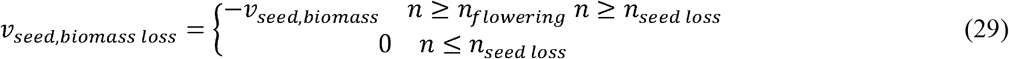

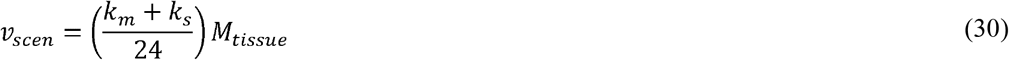

Where *I* is the set of metabolites; *J* is the set of reactions; *Micro* is the set of micronutrients (phosphate, ammonium, and sulfate) and is a subset of *I; X* is the set of amino acids which are synthesized in the leaf tissue and exported to other tissues; and *Y* is the set of sugars stored by various plant tissues. In addition, *M* is defined as a very large number, t_leaf_ is the mass-normalized transpiration rate from the leaf; *s_la_*, *s_sa1_*, and *s_sa2_* are the amplitudes of the sine wave modeling of starch storage in the leaf and stem and sucrose storage in the stem, respectively; *s_lf_*, *s_sf1_*, *s_sf2_* are the frequencies of the sine wave modeling of starch storage in the leaf and stem and sucrose storage in the stem, respectively; *s_lx_*, *s_sx1_*, and *s_sx2_* are the x-intercept shifts of the sine wave modeling of starch storage in the leaf and stem and sucrose storage in the stem, respectively; *s* is the level of seeding of the model; *x_tissue_* is the fraction of total plant mass accounted for by that tissue; and *M_tissue_* is the mass of the given tissue. The following subsections will explain the constraints used in the FBA framework.

### Defining model objective functions

For most analyses and results, the objective function of p-ath780 has been to maximize the sum of the biomass production rates for all four tissues according to the following equation (referred to as the default objective).

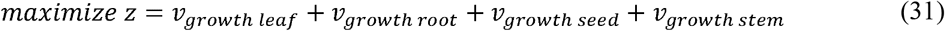

Where *z* has been defined the objective function and *ν_growth tissue_* is defined as the rate of biomass production, in units of h^−1^, of the tissue referenced. This objective function is approximately equivalent to having maximized the growth rate (change in mass per unit time) of the plant as a whole. This objective function has led to one major issue, namely how to avoid the model producing only the metabolically “cheapest” tissue which could result in the maximum objective value but is biologically unrealistic. This is addressed by equations (23) through (28) and will be further discussed later.

It has been noted that the maximization of plant biomass has not been the only feasible objective function for plant SM system, for instance one alternate objective function is the maximization of plant photonic efficiency [15,16]. This objective has generally been framed as minimizing the amount of light used by the plant system, given a required growth rate [15,16]. As it has been assumed that the only (significant) photosynthetic tissue in the p-ath780 model is the leaf tissue, only the objective with relation to the leaf tissue has been altered. As a result, the leaf tissue term in equation (1) has been replaced with a photonic efficiency term in the following equation.

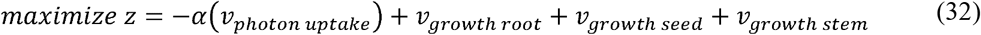

Where *α* has been defined a correction factor to scale *ν_photon uptake_* to be on the same order of magnitude as the growth rates for each other tissue.

An alternative objective function has also been defined for the seed tissue. Specifically, as fatty acids have been shown to one of the most prominent forms of carbon storage in the seed tissue [38], the alternate objective function is the maximization of seed fatty acid stores. This has resulted in seed objective function as follows:

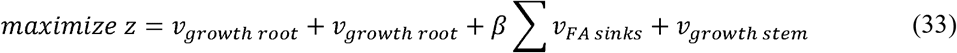

Where the new seed flux term is defined as the sum of fatty acid storing (sink) reactions and *β* serves to reduce this term to be equal in order of magnitude to the other objectives. Similar to *α, β* has been determined through trial and error. One additional objective function has been studied which combines linear photonic efficiency and fatty acid storage:

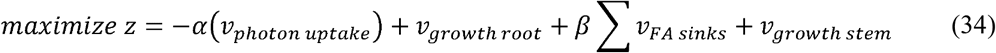

Equation (33) has combined three separate objective types: linear photonic efficiency (leaf), biomass production (root and stem), and fatty acid storage (seed). It should be noted that the objective for the root and stem tissues are always to maximize biomass production. For more details see Supplemental File 22.

### Mass balance

In this model, the mass balance, equation (2), is allowed some flexibility for the storage of metabolites in the plant tissue up to 10 mmol per gDW hour. This has been found to be necessary in the design-build-test cycle so that all points would be feasible.

### Enforcing net CO_2_ consumption O_2_ production

In equations (3) and (4), it is required that the net effect plant metabolism is carbon fixation and oxygen production, since this is a well-known role of plant systems (see Supplemental File 22).

### Enforcing logical flow of micronutrients

In equations (5) through (9) the logical flow of micronutrients is dictated. Equation (5) ensures that the uptake rate is bounded, equation (6) ensures that the rate of each micronutrient exported by the root to the other tissues is less than or equal to that uptaken by the root from the soil, allowing for the root to use a portion of the uptake nutrients. Equation (7) ensures that all micronutrient exported by the root is uptaken by the stem, and equation (8) is essentially the same as equation (6), but for the stem tissue. Finally, equation (9) ensures that the micronutrients exported by the stem are all given to other tissues, specifically leaf or seed. Both sides of the equation in equations (6) through (9) are multiplied by each tissues mass to convert the units of the constraint from mmol per gDW hour to mmol per hour as each tissue has a different mass value in gDW. For more details see Supplemental File 22.

### Enforcing logical flow of amino acids

Similar to micronutrients, the logical flow of amino acids has been defined explicitly via equations (10) to (12), as having been synthesized in the leaf tissue and exported to seed tissue. This is because seed tissue has not been shown to produce all needed amino acids in the MFA study consulted [27], and the root and stem models do not require amino acids for biomass production in the defined biomass composition. Essentially, these constraints ensure that all amino acids exported by the leaf are uptaken by the stem [equation (6)]; that these amino acids are not stored in the stem [equation (7)]; and that all amino acids are exported by the stem to the seed tissue. For more details see Supplemental File 22.

### Enforcing logical flow of sucrose

As with amino acids, sucrose is modeled as being produced in the leaf tissue and exported to other tissues. In contrast to amino acids, sucrose is necessary for all tissue models. Therefore, equation (13) is analogous to equation (10), equation (14) allows the stem to use the sucrose it receives from the leaf, unlike equation (11), and equation (15) exports sucrose both to the seed and root, unlike equation (12). For more details see Supplemental File 22.

### Enforcing diurnal patterns of carbohydrate storage

Plants store carbohydrates in leaf and stem tissues in the form of starch (leaf and stem) and sucrose (stem) in a pattern where the rates of storage may be modeled by a sine wave with a period of 24 hours [31,41]. The calculations for defining the necessary parameters, parameters s_la_, s_sa1_, s_sa2_, s_lf_, s_sf1_, s_sf2_, s_lx_, s_sx1_, and s_sx2_ in equations (16) through (18), of the fit sine waves (see Supplemental Files 5 and 22).

### Enforcing logical flow of water

The flow of water in the p-ath780 model is constrained as similar to that of micronutrients and are defined in equations (19) through (22), but without the equivalents of equations (6) and (8), and with the addition of a transpiration constraint. This difference is because oxidative phosphorylation in tissues creates water. Hence, in tissues without significant photosynthetic activity, water might be produced in this model, and the largest usage to which plants put water, to maintain turgor pressure, is not modeled as SMs use gDW as a basis of calculation, rather than fresh weight (see Supplemental File 22).

### Defining the relationship between tissue growth rates

To avoid the aforementioned problem of having p-ath780 produce only the “cheapest” biomass, the growth rates of all four tissues have been linked by a series of constraints which ensure that they grow at rates which maintain the desired tissue mass ratios. The rate of biomass production determined by a SM is the exponential growth rate of the biological system being modeled [8]; therefore, plant mass can be defined as:

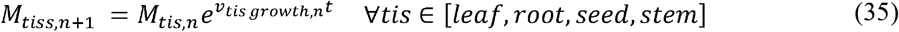

Where *M* has been defined as the plant mass at time *n* + 1, *ν_plant growth, n_* is defined as the rate of plant growth at time *n*, and *M*_*n* – 1_ is defined as the plant mass at time *n*. Further, the ratio of the masses to two tissues can be defined with reference to a single tissue, such as leaf, in the following manner:

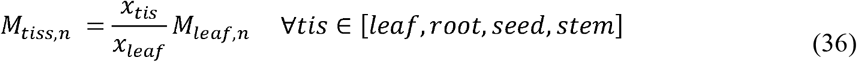

By having substituted the former equation into the latter and simplifying the result (see Supplemental File 22), linear equations have been written to constrain biomass production rates of root, seed, and stem tissues with respect to leaf tissue as follows:

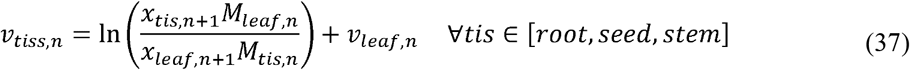

This constraint we have added to the SM model, as equations (23) through (28) in order to ensure that all tissues do have biomass production (or loss) and that it is in an amount which will result in tissue masses in the correct proportions.

### Ensuring non-productive loss of seed mass in silique shattering

A constraint has been found necessary to enforce that metabolites associated with the loss of seed biomass, modeled by the biomass production constraint having reversed flow, when seeds are being lost during silique shattering (in the Silique Ripening Stage) and are not recycled into other parts of plant metabolism. This constraint just does that by forcing recovered metabolites into the biomass loss reaction of the seed tissue.

### Defining model maintenance and senescence costs

An important consideration in any SM is the definition of a maintenance cost, which is typically defined as ATP hydrolysis [26]. Biomass-based maintenance and senescence costs have been defined as they have been suggested as more accurate or applicable for plant systems [36,42], but have not yet been used in an SM. We have defined maintenance and senescence costs as a biomass drain on each tissue scaled by tissue mass in equation (30). A maintenance cost value of k_m_=0.03 day^−1^ has been defined which is in an order of magnitude typical for plant systems [42], and the same value has been defined for plant senescence, k_s_, as this parameter appears to generally be of the same order of magnitude [36,42]. These rates are then converted into their per hour equivalent and scaled by tissue mass to enforce these constraints. Only a single constraint has been defined for both phenomena as both are biomass drains whose effect is additive. Literature evidence, including pictorial evidence of plant phenotype at various growth stages, appears to suggest that the rate of plant senescence increases drastically as the flowering production stage finishes and the Silique Ripening phases begin (in literature, growth stage 0.65 to 9.70) [24]. Further, it appears that the plant no longer maintains current mass, but allows tissues to die and desiccate [24]. This has been included in the p-ath780 model in that plant senescence is increased by two orders of magnitude and plant maintenance is set to zero following the end of the Flower Production stage. This results in a growth curve in-line with *in vivo* evidence (see Table 1).

### Other constraints enforced on the FBA-like optimization framework

There are other constraints enforced on the optimization framework discussed above that are more difficult or cumbersome to state mathematically and are therefore discussed here.

### Defining the usage of seed stores by the seedling

A seedling’s source of carbon is primarily its reserves of stored carbohydrates, proteins, and lipids. Namely, it has been shown that seeds have stores of approximately 0.425 *μ*g of sucrose, 6 *μ*g of fatty acids, and 6 *μ*g of proteins (modeled here as component amino acids) available [30]. As no information concerning the pattern of usage of the seed storage has been found, it has been assumed that the stores are utilized at a constant rate during the duration of the seed germination period and that all the storage is fully consumed by 88.5 hours after germination, which has been defined the point at which the cotyledons are fully open and leaf development intensifies [24]. The rate at which the seedling should have uptaken the seed storage has been determined by identifying the moles (mmol) of each major component of the seed storage and dividing by the time over which the seedling consumes those. This has resulted in a mmol/h quantity. See Supplemental File 5 for this calculation. This quantity has then been scaled by plant mass to result in a mmol/gDW□h quantity, which is used to bound the uptake rates of seed store metabolites. As the leaf has proven the most metabolically active tissue, it is assumed that the leaf tissue of an arabidopsis seedling uptakes the stored fatty acids, amino acids, and carbohydrates which is provided for seedling growth during the Seed Germination stage when the leaves have no access to light (see Figure 1, Seed Germination).

### Defining initial plant and tissue ratios

As the model advances plant and tissue masses with respect to time, the establishment of initial mass for plant and tissues has become important in this framework. Experimental evidence has shown that arabidopsis seeds have a fresh weight (FW) of 25.3 *μ*g and have only about 7% water content [30]. The embryo itself is assumed equal to the seed mass less the mass of seed stores of sucrose (0.425 *μ*g), Fatty Acids (6 *μ*g), and proteins (6 *μ*g) [30]. Having assumed that the dry matter content ratio holds for the embryo as well, this has left approximately 11.0 *μ*g dry weight (DW) for the embryo. As information on the ratio of tissue masses in arabidopsis has not been documented in literature, the general ratio for herbaceous plants has been used as a starting point, namely 0.46:0.24:0.3 leaf:root:stem FW [32]. This ratio has been converted to DW ratio for stoichiometric modeling. Experimental data has shown that the dry matter content of leaf tissue is 0.212 DW/FW, of root tissue is 0.170 DW/FW, and of the stem tissue is 0.176 DW/FW [44]. Having converted the FW ratios to DW ratios has given the ratio of 0.511:0.267:0.211 leaf:root:stem DW. While the dry matter content of an embryonic arabidopsis is much higher than that of a mature plant (the source of the utilized dry matter content ratios), this DW tissue ratio has non-the-less been assumed to be accurate for the embryo due to lack of evidence to the contrary.

### Defining stage times

Time points which define the transition between different stages of growth have been taken from a single source of experimental evidence [24]. Stage transitions selected include the transition to stage 0.70 (Seed Germination to Leaf Development transition in Figure 3), stage 6.00 (Leaf Development to Flower Production transition in Figure 3), and stage 8.00 (Flower Production to Silique Ripening transition in Figure 3). Not all lifecycle stage transitions for which there is experimental evidence have been incorporated into this model. In some cases, this has been due to a lack of metabolic relevance, such as the transition from stage 1.04 to stage 1.05 where the plant transitions from 4 rosette leaves to 5 rosette leaves that are greater than 1mm in length. This has not been important to the p-ath780 model as a ratio of plant mass to leaf surface area ratio is used instead [33] (see Supplemental File 5). Others cannot be modeled by the current framework tissues such as stage 5.10 which is when the first flower bud is visible [24], as the current p-ath780 model has no flower bud tissue. The length of the seed ripening stage is also determined by experimental evidence [24].

### Defining the change in tissue mass ratios with growth stage

Using available literature evidence, two endpoints for the plant tissue mass ratios have been defined when no seeds are present and all seeds are produced [24,38]. The transition between these states are assumed to be linear with respect to a parameter called seeding, defined above as *s*. These relationships are then modeled as:

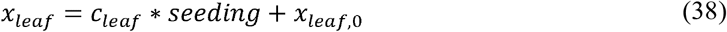

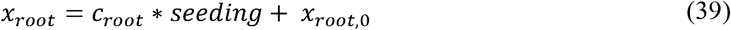

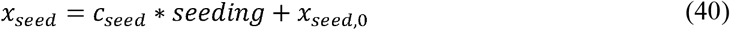

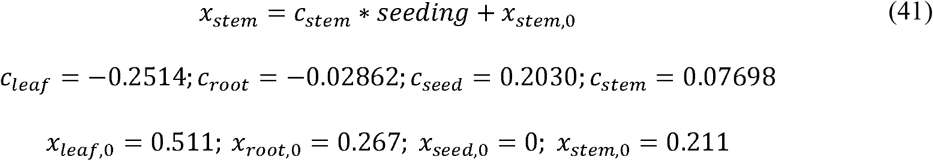

Where *x_tissue_* has been defined as the tissue mass fraction with respect to the total mass of the plant, *c_tissue_* is defined as the change in tissue mass fraction with respect to seeding, and *x_tissue_* is defined as the initial mass fraction of each tissue. The gain in the seeding parameter has been assumed to be linear with time and is fit to experimental time point describing the fraction of flowers produced [24] (see Supplemental Files 5 and 22).

### Defining the availability of light

The amount of light available to the model to use for photosynthesis has been defined initially by literature sources used for other constraints [31], and scaled by the transmittance of that light source (fluorescent lights) [34] and the absorbance of arabidopsis leaves [34] and surface area to plant mass of arabidopsis leaves [33] to define the amount of light usable by the plant system, which has been approximately estimated to be 4.00 mmol/gDW plant·h. This value has been shown to be 21.50% of the total photons output by the fluorescent light (see Supplemental Files 5 and 22).

### Defining the FVA for the p-ath780 model

A Flux Variability Analysis (FVA) model has been defined for growth both in light and dark growth. All flux bounds and constraints are the same and the FBA models, but the objective function is defined as:

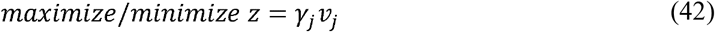

Where the FVA model solution has been iterated for each reaction *j*, and *γ_j_* has been valued at 1 for the current reaction whose maximum and minimum are to be investigated and 0 for all others and is stepped through first maximizing and then minimizing each reaction. Due to restrictions of the time allowed for model solutions, nine points has been selected at which to perform FVA. These points are 70 hours after germination (HAG, seed germination stage, dark), 408 HAG (leaf development stage, light), 576 HAG (leaf development stage, light), 590 HAG (leaf development stage, dark), 800 HAG (flower production stage, light), 810 HAG (flower production stage, dark), 1156 HAG (flower production to silique ripening transition), 1200 HAG (silique ripening stage, light), and 1220 HAG (silique ripening stage, dark).

### Defining the mass step between time points

Using the biomass production rates calculated by the FBA-like optimization framework, a Constrained Non-linear System (CNS) of equations can be defined to advance the plant mass by treating the growth rates as constants. This system of equations has been derived from the basic principles of FBA (e.g. that growth rates are exponential rates of growth) through a sequence of simplifications and assumptions which can be found in Supplemental File 22, and therefore will not be elaborated on here. The end result is shown below for a given time point *t*.

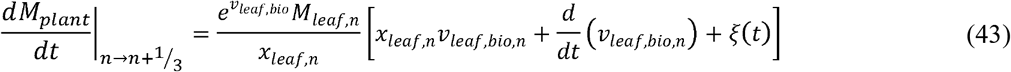

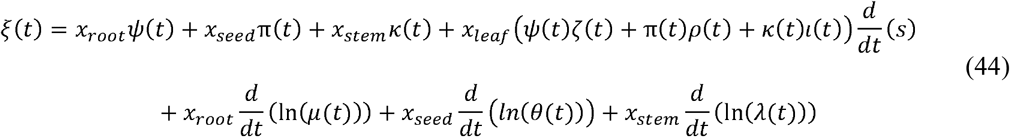

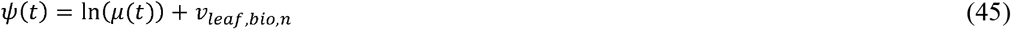

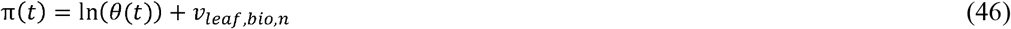

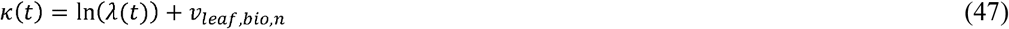

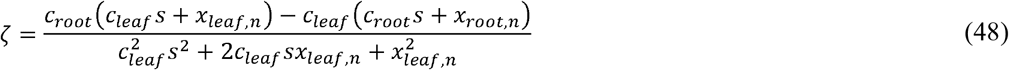

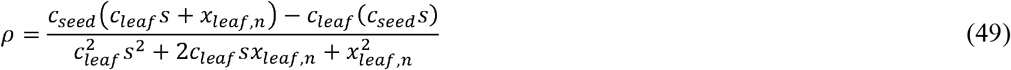

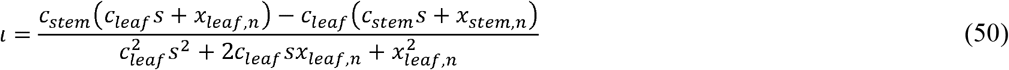

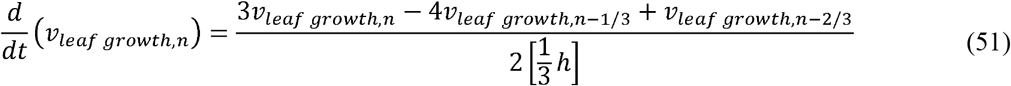

Where, μ, θ, λ are parameters defined in equations (24), (26), and (28), The above system of nine equations has nine corresponding variables: the mass step [LHS of equation (43)], *ξ, ψ, π, ζ, ρ, ι*, and the time derivative of the leaf growth rate [LHS of equation (53)].

Equation (51), as shown above, comes from a backwards finite difference formula of error order *h*^2^. For the purposes of increased model accuracy and stability, the FBA-like framework is solved every ⅓ of an hour to more accurately calculate the mass step at each hour (error is approximately one ninth of that when using full hour values as estimates). As equation (51) is an estimate (rather than an exact equation), should CNS not find a feasible solution, it is further relaxed using a tolerance parameter (*tol*) to a pair of inequalities:

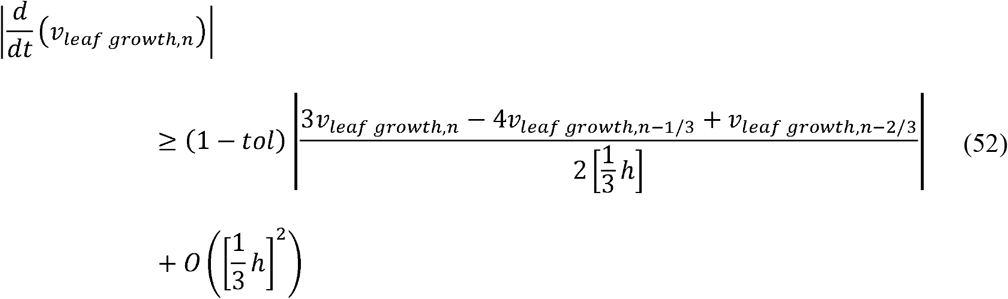

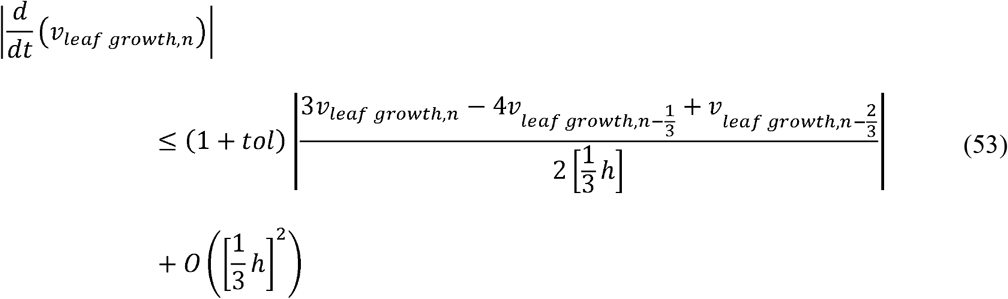

Where, *tol* begins at 0.00 and increases by 0.10 for each iteration if a solution is not found. This results in an mixed-integer non-linear programming (MINLP) problem (8 equality constraints and 9 variables), and the BARON solver has been used to attempt to solve the model. In all cases where a solution has not been found via a CNS solver, a solution has been found using the MINLP solver at *tol* = 0.10. The above set of equations [either equations (43) through (51) as a CNS problem or (43) through (50), (52), and (53)] is solved three times to make estimates of the LHS of equation (42) usable in Heunn’s rule for explicit third-order Runge-Kutta method. For why this method has been used (see Supplemental File 22).

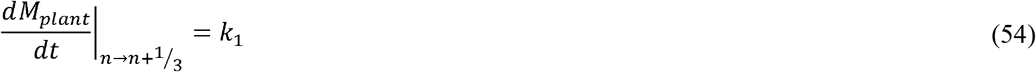

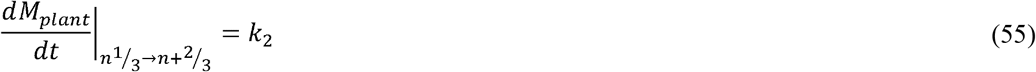

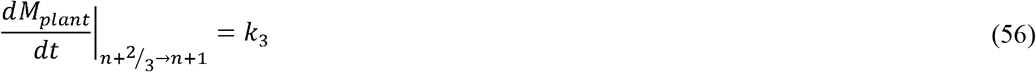

After each partial step, the plant and tissues masses are updated for the next solution. These mass step estimates are then combined using Heunn’s rule for explicit third-order Runge-Kutta method, where the new mass is calculated as follows.

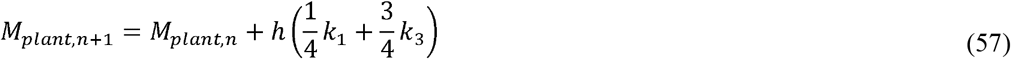

And the mass of each individual tissue is then updated as follows:

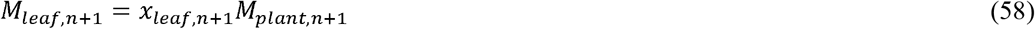

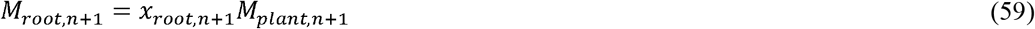

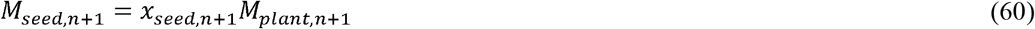

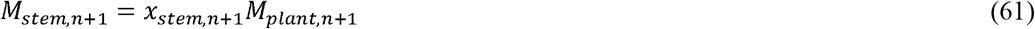

Why p-ath780 updates overall plant mass rather than solving the above problem for each individual tissue is discussed in Supplemental File 22.

## Software platforms used

See Supplemental file 23 for which programming language various supplemental files utilize. For Python code, version 3.3 is used; for Perl, version 5.26 for Supplemental Files 18 through 21 and Strawberry Perl version 5.24.0.1 is used for Supplemental File 8; GAMS code utilizes version 24.7.4. All GAMS and Python code, in addition code included in Supplemental File 8 is run using the Holland Computing Center at the University of Nebraska, Lincoln. Supplemental Files 18 through 21 utilize the additional module the LWP (the world-wide web library for Perl) module 6.39, and have been run on a windows desktop computer.

## Code availability

The authors declare that the code supporting the findings of this study is available within the article’s Supplementary Information files.

## Abbreviations used

For the convenience of our readers, a list of abbreviations used is given below:

GPR: Gene-Protein-Reaction
SM: Stoichiometric Model
FBA: Flux Balance Analysis
FVA: Flux Variability Analysis
LP: Linear Problem
CNS: Constrained Non-linear System
MINLP: Mixed Integer Non-Linear Problem
arabidopsis: *Arabidopsis thaliana*
LHS: Left-Hand Side
wrt: with respect to
gDW: grams Dry Weight
DW: Dry Weight
gFW: grams Fresh Weight
FW: Fresh Weight
MFA: Metabolic Flux Analysis
KEGG: Kyoto Encyclopedia of Genes and Genomes
DAG: Days After Germination
HAG: Hours After Germination

## Acknowledgement

This work has been completed utilizing the Holland Computing Center of the University of Nebraska, which receives support from the Nebraska Research Initiative. The authors gratefully acknowledge funding from UNL Faculty Startup Grant 21-1106-4038.

## Author contributions

Experiments have been conceived by R.S. and W.L.S. W.L.S. performed the experiments and analyzed the data. R.S. and W.L.S. contributed analysis tools. R.S. and W.L.S. wrote the manuscript.

## Supplementary information

is provided. To help readers navigate the extensive set of included data and replicate this study, Supplemental File 23 provides an overview of the included supplemental files and lays out the file structure to use in conjunction with the p-ath780 model.

## Supporting Information Captions

**Supplemental_File_1.txt:** This file is a text file (extension “.txt”) which contains the seed tissue model of p-ath780. This file is referenced as “p-ath780Seed.txt” by other supplemental files, particularly code, and this correct name should replace the default name for attached code to run properly.

**Supplemental_File_2.txt:** This file is a text file which contains the leaf tissue model of p-ath780. This file is referenced as “p-ath780Leaf.txt” by other supplemental files, particularly code, and this correct name should replace the default name for attached code to run properly.

**Supplemental_File_3.txt:** This file is a text file which contains the root tissue model of p-ath780. This file is referenced as “p-ath780Root.txt” by other supplemental files, particularly code, and this correct name should replace the default name for attached code to run properly.

**Supplemental_File_4.txt:** This file is a text file which contains the stem tissue model of p-ath780. This file is referenced as “p-ath780Stem.txt” by other supplemental files, particularly code, and this correct name should replace the default name for attached code to run properly.

**Supplemental_File_5.xlsx:** This file is a Microsoft Excel file which store a wide variety of information concerning the p-ath780 model. This include the manually-curated GPR results for each tissue model, the calculations pertaining to the determination of the biomass equation for each tissue model, calculations for various parameters used in the p-ath780 model to incorporate literature data, and calculations pertaining to the diurnal storage and uptake of carbohydrates.

**Supplemental_File_6.txt:** This file is the GAMS code for the p-ath780 model itself. It is generally named “p-ath780.gms”.

**Supplemental_File_7.txt:** This is an executable Python code which takes the input of a model (such as Supplemental Files 1 through 4) and outputs a number of files which can be read by GAMs code. Generally, this code is named “convert.py”. This code requires slight modifications depending on which model file is to be converted (see in-code comments).

**Supplemental_File_8.txt:** This is an executable Perl code which takes the results of converting each tissue model file using Supplemental File 11 and creates some of the necessary inputs for the p-ath780 GAMS code. This is generally referenced as “makeGrowthInputs.pl”.

**Supplemental_File_9.txt:** This is a text file which contains a list of the names of parameters used to defined p-ath780 model growth. This is referred to by other files as “growthSpecsNames.txt”.

**Supplemental_File_10.txt:** This is a text file which contains the actual specifications used for growing by the p-ath780 model. This is referred to by other files as “growthSpecs.txt”, importantly it is referred to as this by Supplemental File 6.

**Supplemental_File_11.txt:** This is a text file containing the list of time points to iterate over for each day, e.g. this contains each hour of the day, beginning at 0 and ending at 23. This file is referenced by others as “timepointsH.txt”, importantly it is referred to as this by Supplemental File 6.

**Supplemental_File_12.txt:** This is a text file containing a list of days to solve the model over, in this case from day 0 to day 61. This file is referred to by others as “timepoints.txt”, importantly it is referred to as this by Supplemental File 6.

**Supplemental_File_13.txt:** This is a text file containing a list of data labels for much of the data saved at each time point (combination of day and hour) and reported on in the troubleshooting file. This file is referred to by others as “timeData.txt”, importantly it is referred to as this by Supplemental File 6.

**Supplemental_File_14.txt:** This is a text file which lists the time at which the sun rises (or light is made available) each day. At present, light is made available at a default time of 0. This file is referred to by others as “sunrise.txt”, importantly it is referred to as this by Supplemental File 6.

**Supplemental_File_15.txt:** This is a text file which lists the time at which the sun sets (or light is no longer made available) each day. At present, light is made available at a default time of 0. This file is referred to by others as “sunset.txt”, importantly it is referred to as this by Supplemental File 6.

**Supplemental_File_16.txt:** This file basically converts the set time of day to a parameter of equal value. Necessary because mathematical operations cannot be performed on sets. This file is referred to by other files as “timeofday.txt”, importantly it is referred to as this by Supplemental File 6.

**Supplemental_File_17.xlsx:** This is a Microsoft Excel file which contains the results of the p-ath780 model for various alternative objective functions. The sheet tabs indicate which alternative objective function the data corresponds to. The key is as follows:

lpe_g_g_g_X: linear photonic efficiency objective for the leaf, growth objective for other tissues. The number which replaces “X” indicates the run number.

g_g_g_g: Growth objective for all tissues.

nlpe_X: Non-linear photonic efficiency objective, the number which replaces the “X” denotes the run number.

g_g_fa_g_X: Fatty acid storage objective for the seed tissue, growth objective for all other, the number which replaces the “X” denotes the run number.

**Supplemental_File_18.txt:** This is an executable Perl code file which is used to automatically curate the Gene-Protein-Reaction (GPR) links for all tissue models using the KEGG API (advanced programming interface, rest.kegg.jp). Inside the documentation of the code is the instructions for adapting it to investigate the GPR links for each tissue. Generally, this file is named “RxnstoGenes.pl”. This file requires the LWP Perl package.

**Supplemental_File_19.tex:** This file is a comma separated values file. This file contains a list of 73 KEGG pathways for which to get the list of associated reactions. This file is referred to by code as “.csv”, and must have the proper name for the code to function properly. Generally referred to as “PathRxns.csv”.

**Supplemental_File_20.txt:** This file is an executable Perl code file (extension of “.pl”) which automatically generates the lists of reactions associated with various KEGG pathways. The KEGG pathways used are listed in Supplemental File 19. This file is generally named “PathGetRxnsComps.pl”.

**Supplemental_File_21.txt:** This file is an executable Perl code file which is used to automatically read the files created from Supplemental Files 7 to give counts of how many reactions a model has which belong to each of the pathways indicated by Supplemental File 7. Generally, this file is named “ModelPathComp.pl”.

**Supplemental_File_22.docx:** This file is a Microsoft Word file which contains all the calculations simplifications, and rational used for the determination of the function for whole-plant mass step with respect to time.

**Supplemental_File_23.docx:** A Microsoft Word file designed to help navigate other files provided as well as to outline the general file structure and barebones workflow used with the p-ath780 model to make model implementation easier.

