## Supplemental File 7 for "A Computational Framework to Study the Primary Lifecycle Metabolism of *Arabidopsis thaliana*"

# converts reaction list and descrition in a text file to GAMS input format
__author__      = "Mohammad Mazharul Islam and Wheaton Schroeder"
#version 1.0: Mohammad
#version 1.1: Wheaton
#updates associated with version 1.1:
#*updated the output to write all files necessary for fba.gms (added exchange_reactions.txt'
#	irreversible_reactions.txt, and reversible_reactions_no_exchange such that this programs
#	output is the input to fba.gms.
#	This program is dependent on the nomenclature in kathKeys.txt, and was originally written
#	to streamline the time to FBA using personal notation
#*minor streamlining of some lines of code, variable addition associated with previous bullet update
#*added command line output so that the user knows the program ran successfully

#importing the string package
import string
#import the regular expression package
import re

#this is the input file, 'r' denotes read operation
input1=open('p-ath780Seed.txt','r')

#these are the output file names, 'w' denotes write operation
output1=open('metabolites.txt','w')
output2=open('rxntype.txt','w')
output3=open('S_matrix.txt','w')
output4=open('reactions.txt','w')
output5=open('exchange_reactions.txt','w')
output6=open('irreversible_reactions_f.txt','w')
output9=open('irreversible_reactions_b.txt','w')
output7=open('reversible_reactions_no_exchange.txt','w')
output8=open('regulated_reactions.txt','w')


#reading the input file
alllines1=input1.readlines()

#declaring empty lists for metabolites and reactions (where each metabolite and reaction will be added one by one after reading the input file)
#complete metabolite list
met_list=[]
#complete reaction list
rxn_list=[]
#list of irreversible reactions
irr_rxn_list=[]
#list of reactions that are reversible, but not exchange
rev_rxn_no_ex_list=[]
#list of exchange reactions
ex_rxn_list=[]
#list of regulated reactions
reg_rxn_list=[]
#list of sink reactions
sink_rxn_list=[]
#objective species
objective=""
#list of transport reactions
trans_rxn_list=[]

output1.write('/\n') #writing a '/' mark at the beginning of the output file 1, then new line (metabolites.txt)
output2.write('/\n') #writing a '/' mark at the beginning of the output file 2, then new line (rxntype.txt)
output3.write('/\n') #writing a '/' mark at the beginning of the output file 3, then new line (S_matrix.txt)
output4.write('/\n') #writing a '/' mark at the beginning of the output file 4, then new line (reactions.txt)
output5.write('/\n') #writing a '/' mark at the beginning of the output file 5, then new line (exchange_reactions.txt)
output6.write('/\n') #writing a '/' mark at the beginning of the output file 6, then new line (irreversible_reactions_f.txt)
output7.write('/\n') #writing a '/' mark at the beginning of the output file 7, then new line (reversible_reactions_no_exchange.txt)
output8.write('/\n') #writing a '/' mark at the beginning of the output file 8, then new line (regulated_reactions.txt)
output9.write('/\n') #writing a '/' mark at the beginning of the output file 8, then new line (regulated_reactions_b.txt)

for line in alllines1:	#for each line in the input file
	print(line)
	line=line.rstrip()	# stripping off any empty characters at the end of the line
	rxn,des=line.split('\t')	#splitting the remaining line with a tab separator, in this case rxn will give reaction name, and des will give you the rest
	
	#Quick key:
	#rxn - reaction name/number/id/etc
	#des - reaction descrition, the written out chemical reaction, possibly with stoichiometry
	
	#block of text to handle regulated reactions before going into if/else scenarios
	if bool(re.search('reg$',rxn)):	#if has reg tag, note that regular expression search converted to boolean 
		if rxn not in reg_rxn_list:	#if not already it regulated reactions list
			reg_rxn_list.append(rxn)	#append it to list
			output8.write("'"+rxn+"'\n")	#write it to appropriate file
	if bool(re.search('t$',rxn)):	#if has reg tag, note that regular expression search converted to boolean 
		if rxn not in reg_rxn_list:	#if not already it regulated reactions list
			trans_rxn_list.append(rxn)	#append it to list
	if '->' in line and '<->' not in line:	#if a forward reaction	
		reactant=des.split('->')[0] #splitting the "des" at '->' sign, [0] gives the first portion
		product=des.split('->')[1] #[1] gives the second
		dir='->'	#directionality
		state='1'
		#find out if objective function, save objective species
		if bool(re.search("biomass0",rxn)):
			objective=reactant
		if rxn not in irr_rxn_list:
			irr_rxn_list.append(rxn)	#add this reaction to irreversible list
			output6.write("'"+rxn+"'\n")	#writes reaction to irreversible reaction list
				#find out if exchange or not
		if bool(re.search('ex$',rxn)):	#if exchange reaction, not that it is converted to a boolean of a regular expression search
			if rxn not in ex_rxn_list:
				ex_rxn_list.append(rxn)	#add to exchange reaction list
				output5.write("'"+rxn+"'\n") #write reaction to exchange reaction list
		if bool(re.search('si$',rxn)):
			if rxn not in sink_rxn_list:
				sink_rxn_list.append(rxn)
	elif '<-' in line and '<->' not in line:	#if a reverse reaction
		reactant=des.split('<-')[0]
		product=des.split('<-')[1]
		dir='<-'	#directionality
		state='-1'
		if rxn not in irr_rxn_list:
			irr_rxn_list.append(rxn)	#add this reaction to irreversible list
			output9.write("'"+rxn+"'\n")	#writes reaction to irreversible reaction list
				#find out if exchange or not
		if bool(re.search('^EX',rxn)):	#if exchange reaction, not that it is converted to a boolean of a regular expression search
			if rxn not in ex_rxn_list:
				ex_rxn_list.append(rxn)	#add to exchange reaction list
				output5.write("'"+rxn+"'\n") #write reaction to exchange reaction list
		if bool(re.search('si$',rxn)): #find out if sink reaction
			if rxn not in sink_rxn_list:
				sink_rxn_list.append(rxn)
	else:	#if a reversible reaction
		reactant=des.split('<->')[0]
		product=des.split('<->')[1]
		dir='<->'	#directionality
		state='0'
		#find out if exchange or not
		if bool(re.search('^EX',rxn)):	#if exchange reaction, not that it is converted to a boolean of a regular expression search
			if rxn not in ex_rxn_list:
				ex_rxn_list.append(rxn)	#add to exchange reaction list
				output5.write("'"+rxn+"'\n") #write reaction to exchange reaction list
		else:	#otherwise, it is not an exchange reaction
			if rxn not in rev_rxn_no_ex_list:
				rev_rxn_no_ex_list.append(rxn)	#append reversible but not exchange list
				output7.write("'"+rxn+"'\n")	#writes reaction to list of reversible reactions that are not exchange reactions
		if bool(re.search('si$',rxn)): #find out if sink reaction
			if rxn not in sink_rxn_list:
				sink_rxn_list.append(rxn)
	if rxn not in rxn_list:
		rxn_list.append(rxn) 	#if the reaction name was not already in the reaction list, we are adding it
		output2.write( "'"+ rxn + "' " + state)	#writing the output in the rxntype.txt file
		output2.write('\n')	#newline
		output4.write("'"+ rxn + "'")
		output4.write('\n')	#newline
			
	dummy_list=reactant.split(' + ') #splitting the reactant side and getting the individual reactants
	k=0
	while k<len(dummy_list): #for every reactant, splitting the stochiometric coefficients
		ind_react=dummy_list[k]
		ind_react=ind_react.strip()
		if ' ' in ind_react:
			st, react_compt = ind_react.split()
		else:

			st = 1
			react_compt = ind_react
			
	
		if react_compt not in met_list:
			met_list.append(react_compt)  	#if the metabolite name was not already in the metabolite list, we are adding it
			output1.write("'"+react_compt+ "'")
			output1.write('\n')	#newline
		#joining the reactant name and rxn name and stoichiometric coefficinet to write into output3 (Sij.txt)
		stoich_line = "'" + react_compt + "'" +'.'+"'" + rxn +"'" +'  '+'-'+str(st)
		output3.write(stoich_line)
		output3.write('\n')	#newline
				
		k=k+1
			
			
	if product != '':
			
		dummy_list2=product.split(' + ') #splitting the product side and getting the individual products
				
		kk=0
		while kk<len(dummy_list2):  #for every product, splitting the stochiometric coefficients
			ind_prod=dummy_list2[kk]
			ind_prod=ind_prod.strip()
			if ' ' in ind_prod:
#				print(ind_prod)
				st,prod_compt=ind_prod.split()
			else:
				st = 1
				prod_compt = ind_prod
			
			if prod_compt not in met_list:
				met_list.append(prod_compt)
				output1.write("'"+prod_compt+"'")
				output1.write('\n')	#newline

			# joining the product name and rsn name and stoichiometric coefficinet to write into output3 (Sij.txt)
			stoich_line = "'" + prod_compt+ "'" +'.'+"'" +rxn+"'" +'  '+str(st)
			output3.write(stoich_line)
			output3.write('\n')	#newline
				
			kk=kk+1
			
		
			
#add the ending '/' mark to note the end of the file
output1.write('/')
output2.write('/')
output3.write('/')
output4.write('/')
output5.write('/')
output6.write('/')
output7.write('/')
output8.write('/')
output9.write('/')

#outputs to let the user know the stats, and show that something has happened
print("number metabolites: "+str(len(met_list))) 	#printing the total number of metabolites
print("number reactions: "+str(len(rxn_list)))	#printing the total number of reactions
print("irreversible reactions: "+str(len(irr_rxn_list))) #print number of irreversible reactions
print("exchange/sink reactions: "+str(len(ex_rxn_list) + len(sink_rxn_list)))	#print number of exchange reactions
print("objective: "+objective)	#print ojective
print("transport reactions: "+str(len(trans_rxn_list)))
