## Supplemental File 8 for "A Computational Framework to Study the Primary Lifecycle Metabolism of *Arabidopsis thaliana*"

#!usr/bin/perl

#Written by: Wheaton Schroeder
#Written to create input files for p-ath780 model so these don't have to be 
#made by hand. Just making input files that change with the models, where manual
#input is not required. Saves a lot of manual labor in the long run

#NOTES:
#this file is usually named "makeGrowthInputs.pl" in the normal workflow
#This file does first rely on the "convertWLS.py" script being run on each tissue model
#This file assumes that each set of outputs from "convertWLS.py" is contained in its own
#	tissue-specific folder named "p-athTissue"
#Running this script is a prerequisite for running "p-ath780.gms"

use strict;

#read in all the reaction files, remove starting and trailing "/" characters on the files
open(LEAFRXNS, "<p-athLeaf/reactions.txt") or die "could not open leaf reaction list, reason: $!";
chomp(my @leafrxns = <LEAFRXNS>);
shift(@leafrxns);
pop(@leafrxns);

open(ROOTRXNS, "<p-athRoot/reactions.txt") or die "could not open root reaction list, reason: $!";
chomp(my @rootrxns = <ROOTRXNS>);
shift(@rootrxns);
pop(@rootrxns);

open(SEEDRXNS, "<p-athSeed/reactions.txt") or die "could not open seed reaction list, reason: $!";
chomp(my @seedrxns = <SEEDRXNS>);
shift(@seedrxns);
pop(@seedrxns);

open(STEMRXNS, "<p-athStem/reactions.txt") or die "could not open stem reaction list, reason: $!";
chomp(my @stemrxns = <STEMRXNS>);
shift(@stemrxns);
pop(@stemrxns);

#write to output reaction file
open(ALLRXNS, ">all_rxns_1.txt") or die "could not write list of all reactions, reason: $!";

printf ALLRXNS "\/\n";

for (my $a = 0; $a <= $#leafrxns; $a++) {

	printf ALLRXNS "%s\n", $leafrxns[$a];

}

for (my $b = 0; $b <= $#rootrxns; $b++) {

	printf ALLRXNS "%s\n", $rootrxns[$b];

}

for (my $c = 0; $c <= $#seedrxns; $c++) {

	printf ALLRXNS "%s\n", $seedrxns[$c];

}

for (my $d = 0; $d <= $#stemrxns; $d++) {

	printf ALLRXNS "%s\n", $stemrxns[$d];

}

printf ALLRXNS "\/";

#repeat the above procedure for metabolites

#read in all the metabolite files, remove starting and trailing "/" characters on the files
open(LEAFMETS, "<k-athLeaf/metabolites.txt") or die "could not open leaf metabolite list, reason: $!";
chomp(my @leafmets = <LEAFMETS>);
shift(@leafmets);
pop(@leafmets);

open(ROOTMETS, "<k-athRoot/metabolites.txt") or die "could not open root metabolite list, reason: $!";
chomp(my @rootmets = <ROOTMETS>);
shift(@rootmets);
pop(@rootmets);

open(SEEDMETS, "<k-athSeed/metabolites.txt") or die "could not open seed metabolite list, reason: $!";
chomp(my @seedmets = <SEEDMETS>);
shift(@seedmets);
pop(@seedmets);

open(STEMMETS, "<k-athStem/metabolites.txt") or die "could not open stem metabolite list, reason: $!";
chomp(my @stemmets = <STEMMETS>);
shift(@stemmets);
pop(@stemmets);

#write to output metabolite file
open(ALLMETS, ">all_mets.txt") or die "could not write list of all metabolites, reason: $!";

printf ALLMETS "\/\n";

for (my $e = 0; $e <= $#leafmets; $e++) {

	printf ALLMETS "%s\n", $leafmets[$e];

}

for (my $f = 0; $f <= $#rootmets; $f++) {

	printf ALLMETS "%s\n", $rootmets[$f];

}

for (my $g = 0; $g <= $#seedmets; $g++) {

	printf ALLMETS "%s\n", $seedmets[$g];

}

for (my $h = 0; $h <= $#stemmets; $h++) {

	printf ALLMETS "%s\n", $stemmets[$h];

}

printf ALLMETS "\/";
