## Supplemental File 18 for "A Computational Framework to Study the Primary Lifecycle Metabolism of *Arabidopsis thaliana*"

#user/bin/perl -w

#written by: Wheaton Schroeder
#the goal of this program is to take a list of reactions (such as kicked out by ListRxnsCpds.pl) and an organism 
#genome ID (kegg, TXXXXX, were X is any digit) and output a list of reactions with their corresponding genes and 
#enzymes

use strict;
use LWP::UserAgent;
my $UserAgent = LWP::UserAgent->new;
require 'common_functions.pl';

#assign the organism genome code
my $genome = "T00041";

#recall the label of the tissu for which the GPR is being studied
my $tissue_label = "se";

#array to store lines to write
my %writeLine = ( );

#array just for the list of genes, used to determine the number of unique genes
my @genes = ( );

#open and read the file, get the list of reactions
if (! open RXNLIST, "<RxnList.txt") {

	die "Could not open/read RxnList.txt, reason: $!";
	
}

chomp(my @rxns = <RXNLIST>);

#create output file
if (! open GPRTABLE, ">GPR.csv") {

	die "could not create/write GPR.csv, reason: $!";
	
}

#for each reaction:
#1) findlist of EC numbers that catalyze the reaction
#2) map EC numbers to genes using the genome ID given

printf "%s reactions found\n", $#rxns;

for(my $i = 0; $i <= $#rxns; $i++) {

	printf "working on reaction %s\n\n", $rxns[$i];
	my $rxn_temp = $rxns[$i];
	my $tempWriteLine = $rxns[$i];
	
	#saves list of ECs so elements can get added in the correct order
	my $EClist = "(";
	my $geneList = "(";
	
	#clean up the input reaction
	#remove compartment designation
	$rxn_temp =~ s/\[.+\]//g;
	#remove tissue designation
	$rxn_temp =~ s/^$tissue_label//;
	
	#do lookup linking reaction to enzyme
	my $RxntoECurl = "http://rest.kegg.jp/link/enzyme/".$rxn_temp;
	printf "URL: %s\n",$RxntoECurl;
	my $rxntoEC = $UserAgent->get($RxntoECurl);
	$rxntoEC = $rxntoEC->content; 
	
	#if the page is blank (checked by if it has an EC number, the reaction is non-enzymatics, no point going further
	if ($rxntoEC =~ /^\s*$/) {
	
		#no possible gene connection by automated means, go to the next enzyme
		printf "no associated enzyme found, non-enzymatic reaction\n\n";
		$tempWriteLine .= ",non-enzymatic,non-enzymatic";
		$writeLine{$rxns[$i]} = $tempWriteLine;
		next;
		
	} else {
		
		#for each EC number
		while ($rxntoEC =~ /ec:(.+)/g) {
		
			#save the value.
			my $EC = $1;
			
			#no need to add delimiter if first element
			$EClist .= $EC;
			printf "found enxyme classification ".$EC."\n";
			
			chomp($EC);
			
			#get the genes related to that EC for the given genome
			my $ECtoGenesurl = "http://rest.kegg.jp/link/".$genome."/".$EC;
			my $ECtoGene = $UserAgent->get($ECtoGenesurl);
			$ECtoGene = $ECtoGene->content;
			
			#read and save each of the gene names
			while ($ECtoGene =~ /ec:.+?\s+(.+)/g) {
			
				my $gene = $1;
				#add gene to gene list, no need to add delimiter for first element
				push @genes, $gene;
				
				$geneList .= $gene;
				printf "found gene ".$gene."\n";
				
				$geneList .= "|";
				
			}
			
			$geneList =~ s/\|$//;
			
			$geneList .= ")||(";
			$EClist .= "|";
			
		}
	
	}
	
	$EClist =~ s/\|$/\)/;
	$geneList =~ s/\|\|\($//;
	
	$tempWriteLine .= ",".$EClist.",".$geneList;
	$writeLine{$rxns[$i]} = $tempWriteLine;
	printf "\n";
	
}

#get the list of unique genes by making the gene list non-redundant
my $nrRef = &RemoveRedundancies(\@genes);
@genes = @$nrRef;

#write to the output file
select GPRTABLE;

printf "Reaction ID,ECs that can catalyze this reaction,Genes that can produce these enzymes\n";

for (my $k = 0; $k <= $#rxns; $k++) {

	printf "%s\n", $writeLine{$rxns[$k]};
	
}

printf "\n\n";

printf "List of all unique genes which can be accounted for by this model\n";
printf "total number: %s\n", $#genes;

for (my $p = 0; $p <= $#genes; $p++) {

	printf "%s\n", $genes[$p];
	
}

close GPRTABLE;
