## Supplemental File 20 for "A Computational Framework to Study the Primary Lifecycle Metabolism of *Arabidopsis thaliana*"

#usr/bin/perl -w

#Written by: Wheaton Schroeder to get a complete reaction and compounds
#for the 75 pathways checked by Model Seed
#input file is a simple csv column of map ids for each pathway of interest
#commas are optional at the end of each of these cells (pathList.csv)
#the output file give potential reactions that can fit into different orthologies
#where the actual/max number of reactions will be closer to the number of orthologies

use LWP::UserAgent;
require 'common_functions.pl';

#open list of path map IDs
if ( ! open MAPS, "<pathList.csv") {
	die "Could not find/access pathList.csv, reason: $!"; #fatal error
}

#array to store pathway names in some form of order
my @pathwayNames = ( );
#hash to input pathway name and get out a list of reactions in that pathway
my %pathtoRxns = ( );
#has to input pathway name and get out a list of compounds in that pathway
my %pathtoCompds = ( );
#keeps an array with orthologies
my %pathtoOrth = ( );
#orthology hash, to determine the % of orthologies with at least one qualifying reaction
#reaction, return orthologies that are satisfied
my %rxntoOrth = ( );
#orthology hash, which when inverted gives the previous hash
my %orthtoRxn = ( );

#read all the map IDs
chomp(@mapIDs = <MAPS>);

for (my $i = 0; $i <= $#mapIDs; $i++) {
	printf "given map ID: %s\n", $mapIDs[$i];
	#temporary reaction list
	@tempRxnList = ( );
	#temporary compound list
	@tempCompdList = ( );
	#temporary orthology list
	@tempOrthList = ( );
	#temporary pathway name
	$pathName = " ";
	$orthCount = 0;
	
	#logic of this operation mapXXXXX page to module pages (MXXXXX) which give the 
	#included compounds and reactions. Those are pulled and added to the hashes
	$mapIDs[$i] =~ s/,//ig; #remove comma, if present
	$koID = $mapIDs[$i];
	$koID =~ s/map/ko/ig;
	#define url to get to pathway page
	$koURL = "http://rest.kegg.jp/get/".$koID."/";
	#create a new user agent which pulls the KEGG API page
	$UserAgent = LWP::UserAgent->new; #creates new user agent
	$koPage = $UserAgent->get($koURL);
	$koPage = $koPage->content; 
	#get the pathway name
	if ($koPage =~ /NAME\s+(.+)/) {
		#add pathway name to list
		push @pathwayNames, $1;
		#define temporary pathway name
		$pathName = $1;
		printf "found path name: %s\n", $pathName;
	}
	#this doensn't list reactions, just modules within the pathway
	#we need to get the modules. This loops for each module "hit" in a 
	#text search
	while($koPage =~ /\s+K[0-9][0-9][0-9][0-9][0-9]\s+/g) {
		@orthRxns = ( );
		#define the module
		$orthology = $&;
		$orthology =~ s/\s+//ig; #remove all spaces in module ID, chomp doesn't do the trick
		push @tempOrthList,$orthology;
		printf "Orthology: %s Reactions: ", $orthology;
		#go to the module page, get all the information from that
		$orthologyURL = "http://rest.kegg.jp/get/".$orthology;
		$orthologyPage = $UserAgent->get($orthologyURL);
		$orthologyPage = $orthologyPage->content;
		#for each reaction
		while ($orthologyPage =~ /R[0-9][0-9][0-9][0-9][0-9]/g) {
			printf "%s ", $&;
			#add the reaction
			push @tempRxnList, $&;
			push @orthRxns, $&;
			#add reaction compounds to list
			$rxnURL = "http://rest.kegg.jp/get/".$&;
			$rxnPage = $UserAgent->get($rxnURL);
			$rxnPage = $rxnPage->content;
		}
		printf "\n";
		$orthtoRxn{$orthology} = join ",",@orthRxns;
	}
	
	#by this point, all reactions and compounds should be known
	#remove redundancies
	$nrRxnList = &RemoveRedundancies(\@tempRxnList);
	@tempRxnList = @$nrRxnList;
	$nrCompdList = &RemoveRedundancies(\@tempCompdList);
	@tempCompdList = @$nrCompdList;
	#update hashes
	$pathtoRxns{$pathName} = join ",",@tempRxnList;
	$pathtoCompds{$pathName} = join ",",@tempCompdList;
	$pathtoOrth{$pathName} = join ",",@tempOrthList;
	
	#invert orthtoRxn
	#get list of orthologies
	$orthList = $pathtoOrth{$pathName};
	#turn it into an array
	@ListofOrths = split /,/,$orthList;
	printf "orthologies found: %s\n", $orthList;
	#for each orthology
	for(my $k = 0; $k <= $#ListofOrths; $k++) {
		#get a list of reactions in orthology
		$rxnsinOrth = $orthtoRxn{$ListofOrths[$k]};
		printf "reactions in %s: %s\n", $ListofOrths[$k], $rxnsinOrth;
		#turn it into an array
		@listofRxnsinOrth = split /,/,$rxnsinOrth;
		#for each reaction in a given orthology
		for(my $p = 0; $p <= $#listofRxnsinOrth; $p++) {
			#if the reaction entry aready exists, supplement it with the new orthology
			if (exists $rxntoOrth{$listofRxnsinOrth[$p]}) {
				$rxntoOrth{$listofRxnsinOrth[$p]} = $rxntoOrth{$listofRxnsinOrth[$p]}.",".$ListofOrths[$k];
			} else { #if it does not exist, create a new entry with this as the only orthology
				$rxntoOrth{$listofRxnsinOrth[$p]} = $ListofOrths[$k];
			}
		}
	}
}

#check to see if the above inversion worked
foreach $key (sort keys %rxntoOrth) {
	$value = $rxntoOrth{$key};
	printf "%s => %s\n", $key, $value;
}


#open file to write to, one for each pathway
if ( ! open OUTPUTRXN, ">pathRxns.csv") {
	die "Could not write/access pathRxns.csv, reason: $!"; #fatal error
}
select OUTPUTRXN;

#write files for each pathway
for (my $j = 0; $j <= $#pathwayNames; $j++) {
	$rxns = $pathtoRxns{$pathwayNames[$j]};
	#print rows of rections and compounds, include labels
	printf "%s,%s\n", $pathwayNames[$j],$rxns;
}

close OUTPUTRXN;

#open file to write to, one for each pathway
if ( ! open OUTPUTCMD, ">pathCmd.csv") {
	die "Could not write/access pathCmd.csv, reason: $!"; #fatal error
}

select OUTPUTCMD;

#write files for each pathway
for (my $j = 0; $j <= $#pathwayNames; $j++) {
	$compds = $pathtoCompds{$pathwayNames[$j]};
	#print rows of rections and compounds, include labels
	printf "%s,%s\n", $pathwayNames[$j],$compds;
}

close OUTPUTCMD;

#open file to write to, one for each pathway
if ( ! open OUTPUTORTH, ">pathOrth.csv") {
	die "Could not write/access pathOrth.csv, reason: $!"; #fatal error
}

select OUTPUTORTH;

#write files for each pathway
for (my $j = 0; $j <= $#pathwayNames; $j++) {
	$orths = $pathtoOrth{$pathwayNames[$j]};
	#print rows of rections and compounds, include labels
	printf "%s,%s\n", $pathwayNames[$j],$orths;
}

close OUTPUTORTH;
