## Supplemental File 21 for "A Computational Framework to Study the Primary Lifecycle Metabolism of *Arabidopsis thaliana*"

#usr/bin/perl -w

#Written by: Wheaton Schroeder to get a complete reaction and compounds
#for 75 pathways
#input file is a simple csv column of map ids for each pathway of interest
#commas are optional at the end of each of these cells (pathList.csv)

use LWP::UserAgent;
require 'common_functions.pl';

#changing this variable updates all the filenames so that it doesn't have
#be be done manually and helps insure that the results of one tissue do
#not override the results of another tissue.  
my $tissue_code = "st";

#open list of path reaction IDs
if ( ! open PATHRXNS, "<pathRxns.csv") {

	die "Could not find/access pathRxns.csv, reason: $!"; #fatal error

}

#array to store pathway names in some form of order
my @pathwayNames = ( );
#hash to input pathway name and get out a list of reactions in that pathway
my %pathtoRxns = ( );
#has to input pathway name and get out a list of compounds in that pathway
my %pathtoCompds = ( );
#stores fraction of orthologies satisfied, same order as pathwayNames
my @fracSatRxn = ( );
#stores fraction of orthologies satisfied, same order as pathwayNames
my @fracSatCmd = ( );

#read all the reaction IDs, each array element is a single pathway with its reaction list 
chomp(@pathRxnIDs = <PATHRXNS>);

#use the reactions list input file to construct the arrays and hashes
for (my $j = 0; $j <= $#pathRxnIDs; $j++) {

	my @line2 = split /,/, $pathRxnIDs[$j];
	my $path2 = shift @line2;
	
	if (! ($path2 ~~ @pathwayNames)) {
	
		push @pathwayNames, $path2;
		
	}
	
	$pathtoRxns{$path2} = join ",", @line2;
	
}

#now check the completeness of the pathway as measured by fraction of reactions present in said pathway
#open list of path map IDs
if ( ! open RXNIDS, "<".$tissue_code."rxns_path780.txt") {

	die "Could not find/access ".$tissue_code."rxns_path780.txt, reason: $!"; #fatal error
	
}

#now check the completeness of the pathway as measured by fraction of compounds in said pathway
#open list of path map IDs
if ( ! open CMDIDS, "<".$tissue_code."mets_path780.txt") {

	die "Could not find/access ".$tissue_code."mets_path780.txt, reason: $!"; #fatal error
	
}

#read all the reaction IDs
chomp(my @RxnIDs = <RXNIDS>);
#read all the compound IDs
chomp(my @CmdIDs = <CMDIDS>);
#remove redundant entries
$nrRxnIDs = &RemoveRedundancies(\@RxnIDs);
@RxnIDs = @$nrRxnIDs;
$nrCmdIDs = &RemoveRedundancies(\@CmdIDs);
@CmdIDs = @$nrCmdIDs;

#for each pathway, see what fraction of reactions are present in the list
for (my $v = 0; $v <= $#pathwayNames; $v++) {

	$pathRxnCount = 0;
	$pathRxnString = $pathtoRxns{$pathwayNames[$v]};
	@pathRxns = split /,/,$pathRxnString;
	$totalPathRxns = $#pathRxns;
	
	#check each reaction, increment count if reaction present in pathway
	for(my $m = 0; $m <= $#RxnIDs; $m++) {
	
		#if the reaction is in the pathway
		if($RxnIDs[$m] ~~ @pathRxns) {
		
			$pathRxnCount++; #increment the count
			
		} #else not do nothing
		
	}
	
	
	
	$fracSatRxn = $pathRxnCount;
	#push satisfaction value
	push @fracSatRxn, $fracSatRxn;
	
	#do similar for compounds
	$pathCmdCount = 0;
	$pathCmdString = $pathtoCompds{$pathwayNames[$v]};
	@pathCmds = split /,/,$pathCmdString;
	$totalPathCmds = $#pathCmds;
	for (my $m = 0; $m <= $#CmdIDs; $m++) {
	
		if($CmdIDs[$m] ~~ @pathCmds) {
		
			$pathCmdCount++;
			
		}
		
	}
	
	$fracSatCmd = $pathCmdCount;
	push @fracSatCmd, $fracSatCmd;
	
}

#add transport and exchange pathway names
push @pathwayNames, "exchange";
push @pathwayNames, "transport";

my $exrxncount = 0;
my $trxncount = 0;

for(my $a = 0; $a <= $#RxnIDs; $a++) {

	#check if exchange or transport reaction, these will be additional categories
	if ($RxnIDs[$a] =~ /ex$/) {
	
		$exrxncount++;
	
	} elsif ($RxnIDs[$a] =~ /t$/) {
	
		$trxncount++;
	
	} #otherwise not exchange or transport
	
}

push @fracSatRxn, $exrxncount;
push @fracSatRxn, $trxncount;

#open file to write to, one for each pathway
if ( ! open FRACSATRXN, ">".$tissue_code."path780RxnPaths.csv") {

	die "Could not write/access ".$tissue_code."path780RxnPaths.csv, reason: $!"; #fatal error
	
}

select FRACSATRXN;

#write files for each pathway
for (my $j = 0; $j <= $#pathwayNames; $j++) {

	printf "%s,%s\n", $pathwayNames[$j],$fracSatRxn[$j];
	
}

close FRACSATRXN;

#open file to write to, one for each pathway
if ( ! open FRACSATCMD, ">".$tissue_code."path780CompdPaths.csv") {

	die "Could not write/access ".$tissue_code."path780CompdPaths.csv, reason: $!"; #fatal error
	
}

select FRACSATCMD;

#write files for each pathway
for (my $j = 0; $j <= $#pathwayNames; $j++) {

	printf "%s,%s\n", $pathwayNames[$j],$fracSatCmd[$j];
	
}

close FRACSATCMD;
