## Supplemental File 22 for "A Computational Framework to Study the Primary Lifecycle Metabolism of *Arabidopsis thaliana*"

**Supplemental File 22: Supplemental Methods**

Wheaton L. Schroeder and Rajib Saha

### **Definition of biomass composition.**

*With respect to seed biomass.* The MFA study referenced had identified the biomass amino acid composition with respect to 14 amino acids (others destroyed during hydrolysis) and used proteinogenic statistical analysis to identify the relative abundance of the other amino acids in biomass. The biomass composition with respect to lipids and carbohydrates was also determined in this MFA, but no integrated biomass composition (comparing relative abundance of individual amino acids, lipids, and carbohydrates in a single analysis) was provided [26]. Therefore, the MFA-determined rates of fatty acids, amino acids, and carbohydrates being added to plant biomass as sink reactions have been used as a basis of comparison to determine the relative amounts of each type of molecule in seed biomass. See Supplemental File 5 for these calculations.

*With respect to leaf biomass.* In the SM study used to define leaf biomass composition [18], histidine has been added to the biomass equation as it was not present, and certain compounds comprising a very small mass fraction of biomass, such as coniferyl alcohol and hexadecanoic acid, have not been included so as to reduce model size without unduly sacrificing accuracy of biomass composition. See Supplemental File 5 for how leaf biomass was determined.

**Reaction confidence scoring.**

Reactions with score of zero either have been user-defined reactions with no direct physical equivalent, such as the biomass reaction, which are nonetheless essential in stoichiometric modeling or are artificial and blocked reactions that have been defined in order to help organize the model file by KEGG-defined pathways to make manual editing easier. Reactions with a score of one have been determined as reactions necessary for modeling, but for which we have provided no literature-based evidence for their inclusion. These reactions are typically non-enzymatic, or transport/exchange reactions required for biomass production. Reaction confidence scores of two, three, or four indicate that evidence exist for its inclusion in the model. For most reactions, genomic evidence (score of two) has been considered sufficient for their inclusion. Genomic evidence has been automatically determined using Supplemental File 18, which automates the establishment of GPR links using KEGG. This is typically done by identifying proteins by their Enzyme Classification (EC) numbers associated with each reaction, then identifying arabidopsis genes which can produce those enzymes. Some reactions without KEGG identifiers have had their GPR defined in the same manner, but manually as opposed to in an automated workflow. This has included oxidative phosphorylation and photosynthesis. For reactions without defined GPR links, knock-in/knock-out type studies (score of three) were not been consulted. This is because direct literature evidence (score of four) is preferable and can be found for much of core-carbon metabolism as *Arabidopsis thaliana* is a model plant species.

### **Defining p-ath780 objective functions.**

All objective function types (i.e. photonic efficiency, fatty acid production, and biomass production) have been defined as a single objective function as the multi-level optimization framework for implementing each objective separately results in opposing goals. For instance, zero photon uptake would have used the minimum quantity of photons, and therefore would be “high efficiency”, but would often have resulted in no growth. Alternatively, maximum growth would have required a high uptake rate of photons for fixation of carbon necessary to maintain that growth, resulting in low “photonic efficiency” in the classical definition. Through equation (2) (as explained in the main manuscript file) these two objectives can be balanced through $\alpha$. Table 1 shows the differences in plant growth and metabolism that result from this mixed objective function for values of $\alpha$.

**Enforcing the logical flow of metabolites between tissues.**

It has been well known that each tissue in a plant system is capable of gas exchange. It has also been well known that root tissue is generally non-photosynthetic and we have assumed that the carbon fixation (photosynthesis) of the stem and seed tissues are negligible compared to that of the leaves. This is because we have assumed that photosynthetic rate is a function of surface area, and it is well known that the surface area of the stem and seed tissues are much less than that of the leaf tissue. Therefore, root, stem, and seed tissues have been constrained such that they uptake oxygen and export carbon dioxide across the system boundary (dashed line in the system diagrams of Fig. 1). The leaf tissue has been constrained such that the carbon fixation rate when light is available is greater than the rate at which carbon dioxide is exported by the other tissue so that the net effect of the entire plant system is carbon fixation. In all tissue interaction constraints, including that for carbon dioxide and those described later in this sub-section, exchange flux rates (units of mmol/gDW‧h) have been scaled by the mass of each tissue (gDW) to ensure mole balance (all fluxes having units of mmol/h) in these constraints. It has been well known that the roots of a plant uptake micronutrients (such as nitrogen, phosphorous, and sulfur sources), ions (such as protons), and water for use by the rest of the plant system. Similarly, it has been well known that the stem tissue transports these nutrients to other plant tissue. Therefore, in p-ath780, the root tissue has uptaken water, phosphate, ammonia, sulfate, and protons across the system boundary. Tissue interaction constraints are then defined such that the amounts of water, phosphate, ammonia, sulfate, and protons stem has uptaken is less than the amount uptaken by the root. Similarly, the amounts of water, phosphate, ammonia, sulfate, and protons uptaken by leaf and seed tissue combined have been constrained to be less than or equal to that exported by the stem. In addition, it has been well known that during the daylight hours, plants lose water through the process of transpiration. In p-ath780, transpiration is modeled as the export of water from the leaf to outside the system boundary. Transpiration for arabidopsis is measured as 0.2 mmol H_2_O‧m^-2^‧s^-1^ [29]. Scaling this by the leaf surface area ratio of arabidopsis, 0.0398 m^2^ leaf surface area per gDW leaf biomass [28], has given a transpiration rate of 28.6315 mmol/gDW‧h with respect to leaf mass.

Amino acid interactions between tissues have been defined out of modeling necessity. From a Metabolic Flux Analysis (MFA) of arabidopsis seed tissue, 14 amino acids have been identified and their rates of production from pathways inherent to the seed tissue were identified [26]. In truth, 16 amino acids had been measured, but due to the methods used by the study, aspartic acid and asparagine were indistinguishable (listed as one amino acid Asx), as were glutamic acid and glutamine (listed as Glx) [26]. As seed biomass has been determined to be composed of all 20 amino acids [26] and we have found no evidence for the synthesis of the remaining amino acids in the seed tissue, we have assumed that the remaining amino acids were synthesized in the leaf tissue and transported to the seed tissue. A total of 14 amino acids have been allowed to be transported between leaf and seed tissue via the stem tissue, including the four amino acids not produced by the seed tissue: arginine, cysteine, methionine, and tryptophan, and ten other amino acids which the seed tissue model cannot synthesize at a rate equal to its needs for growth or were otherwise found necessary for plant system growth were allowed to be transported between tissues. These ten extra amino acids are: glutamate, glycine, alanine, lysine, aspartate, glutamine, serine, proline, valine, and threonine. All amino acids use the same set of three constraints. The net effect of these constraints has been that the amount of any amino acid exported by the leaf tissue is equal to the amount uptaken by the seed tissue, but that the metabolites are required to first pass through the extracellular environment of the stem tissue (e.g. pass through some vascular tissue).

It has been well known that sugars synthesized in the leaves are transported to non-photosynthetic tissues via plant vascular tissue. In p-ath780, this has been modeled by three constraints which ensure that: i) the amount of sucrose uptaken by the stem is equal to the amount exported by the leaf, ii) the amount of sucrose exported by the stem is less than or equal to that imported by the stem, iii) the amount of sucrose uptaken by the root and seed tissues combined is equal to the amount of sucrose exported by the stem tissue.

### **Defining the diurnal patterns of carbohydrate storage.**

It has been experimentally shown that plant tissues store sugars during periods of growth where light is available, and consume those store when light is no longer available [32]. Experimental data concerning the extent to which this occurs over a 12 hours light and 12 hours dark diurnal metabolic cycle has shown that the peak storage of starch in the leaf tissue is approximately 32.8 $\mu$mol hexose/gFW, while the trough storage is approximately 3.6 $\mu$mol hexose/gFW on a 24-hour period [32]. The slope of this curve has been used to define the rate at which starch is uptaken or stored (see Supplemental File 5). This storage/uptake cycle has been fit to a sine wave with a 24-hour period (frequency 0.261799 h^-1^) with an amplitude of 9.44‧10^-3^ mmol starch dimer/gDW‧h and an x-intercept of 0.0726 hours (indicating the plant takes about 4.3 minutes to switch between storage and usage modes). Similarly, it has been demonstrated that the shoot (stem) tissue of arabidopsis likewise stores and uptakes starch and sucrose [37] in a manner which can be similarly fit to a sine curve to describe rate. These sine curve amplitudes have been determined to be 1.66‧10^-2^ mmol/gDW‧h and 1.25‧10^-3^ mmol/gDW‧h of starch dimer and sucrose, respectively. Both curves have a 24-hour period and an x-intercept of zero. In p-ath780, these sine curves form constraints which have required uptake or storage of starch and sucrose at the appropriate points in the diurnal cycle in all stages where photosynthesis occurs (e.g. all except Seed Germination). In the exceptional case of Seed Germination, internal starch storage and uptake has been constrained to zero.

### **Defining the change in tissue mass ratios with growth stage.**

It has been assumed that the ratios between leaf, root, and stem tissues are consistent with that of a generic herbaceous plant, 0.511:0.267:0.211 leaf:root:stem DW [31], for all growth states. Combining this with an initial seed tissue mass of 0, this forms one end point for tissue mass ratio change modeling for when *s=0*. For stages which have seed mass (Flower Production, Flower Production to Silique Ripening Transition, and Silique Ripening) a parameter, called seeding, defined as *s* above, has been defined which indicates the fraction of maximum seed mass that is currently a part of the plant system. In determining the rate of tissue mass fraction change, the leaf, seed, and stem harvest masses have been noted as 163.7 mg DW, 127.9 mg DW, and 188 mg DW, respectively [24], and this is used as the other end point for tissue mass ratio change when *s=1*. When assuming that the ratio of 0.267:0.211 root:stem tissue holds, when all seeds are produced and part of the plant system, the tissue mass ratios are 0.2598:0.2387:0.2030:0.2984 leaf:root:seed:stem DW. It is assumed that the transition between these states is linear wrt the seeding parameter which can be used to update the tissue mass ratios. These relationships of tissue mass fraction of entire plant DW are defined as follows:

$$x_{leaf}=c_{leaf}*seeding+x_{leaf,0}$$

(36)

$$x_{root}=c_{root}*seeding+ x_{root,0}$$

(37)

$$x_{seed}=c_{seed}*seeding+x_{seed,0}$$

(38)

$$x_{stem}=c_{stem}*seeding+x_{stem,0}$$

(39)

$$c_{leaf}=-0.2514;c_{root}=-0.02862;c_{seed}=0.2030;c_{stem}=0.07698$$

$$x_{leaf,0}=0.511; x_{root,0}=0.267; x_{seed,0}=0; x_{stem,0}=0.211$$

Where $x_{tissue}$ has been defined as the tissue mass fraction with respect to the total mass of the plant, $c_{tissue}$ is defined as the change in tissue mass fraction with respect to seeding, and $x_{tissue}$ is defined as the initial mass fraction of each tissue. The gain in the seeding parameter has been assumed to be linear with time, and is fit to experimental time point describing the fraction of flowers produced [24], see Supplemental File 5 for calculations. Due to the lack of data suitable for modeling purposes and the lack of a flower model in p-ath780, it is assumed that flower tissue is equivalent to seed tissue, and it has been assumed that the fraction of flowers open can be equated to the seeding parameter. For example, when 10% of total flowers have opened at time 35.9 days after germinating (DAG) then seeding has a value of 0.1. Using a linear regression fit, we have determined that the seeding parameter gains 0.00237 per hour during the Flower Production stage. Similarly, as the first seeds are lost at 48 DAG and all seeds have been loosed at 61 DAG, a linear regression has resulted in a seeding parameter loss of -0.00359 per hour in the Silique Ripening stage (Figure 1). The overlap between these two stages has used a per hour seeding parameter change equal to the sum of the change in the Flower Production and Silique Ripening stages (-0.00109 per hour) (Flower Production to Silique Ripening Transition, Figure 1).

### **Defining the availability of light.**

According to a source which we have used for other experimental data which informs p-ath780, arabidopsis plants have grown under 12 hour light and 12 hour dark diurnal cycles under 130 $\mu$mol/m^2^·s intensity fluorescent light [32]. Fluorescent light has thin transmittance peaks at 544 and 609 nm [33], whereas peak absorption for arabidopsis leaves are 440 and 680 nm [34]; therefore, the light which has been provided to the plant for growth does not perfectly match that which can be used by the plant for photosynthesis. By having assumed that all light absorbed by the leaf is usable in photosynthesis (either photosystem), and calculating the number of available photons for the photosystem, it has been determined that the availability of photons for photosystem use is 100.6 mmol/m^2^·s. This has been converted to a usable form for modeling by defining the availability of photons in terms of plant dry weight. Using an average value for leaf surface area per mg DW plant [34], the availability of photons for the photosystem is 4.00 mmol/gDW plant·h. This value has been shown to be 21.50% of the total photons output by the fluorescent light. See Supplemental File 5 for details on this calculation.

### **Defining the mass step between time points.**

Generally, if we could assume a constant exponential growth rate the differential equation for exponential growth can be defined as:

$$\frac{dM}{dt}=v_{growth}M$$

Problem: $v_{growth}$ is also a function of time and plant mass, so this differential equation is not strictly true. We know that exponential growth can be described as:

$$M=M_{0}e^{v_{growth}*t}$$

Taking the time derivative:

$$\frac{dM}{dt}=M_{0}*\frac{d}{dt}\left( e^{v_{growth}*t} \right)$$

Using the chain rule:

$$\frac{dM}{dt}=M_{0}*\frac{d}{du}\left( e^{u} \right)*\frac{du}{dt}$$

$$u=v_{growth}*t$$

$$\frac{du}{dt}=\frac{d}{dt}\left( v_{growth}*t \right)=v_{growth}*\frac{d}{dt}\left( t \right)+t*\frac{d}{dt}\left( v_{growth} \right)=v_{growth}+t*\frac{d}{dt}\left( v_{growth} \right)$$

Where the derivative of $u$ was found using the product rule. Substituting and solving we get:

$$\frac{dM}{dt}=M_{0}*\frac{d}{du}\left( e^{u} \right)*\left( v_{growth}+t*\frac{d}{dt}\left( v_{growth} \right) \right)$$

$$\frac{dM}{dt}=M_{0}*e^{u}*\left( v_{growth}+t*\frac{d}{dt}\left( v_{growth} \right) \right)$$

$$\frac{dM}{dt}=M_{0}*e^{v_{growth}*t}*\left( v_{growth}+t*\frac{d}{dt}\left( v_{growth} \right) \right)$$

Some numerical approximation of $\frac{dv_{growth}}{dt}$ will need to be made. However, we do not know the total growth rate, instead we know the growth rate of individual tissues, therefore more accurately we have:

$$M_{plant}=M_{leaf,0}e^{v_{leaf,bio}*t}+M_{root,0}e^{v_{root,bio}*t}+M_{seed,0}e^{v_{seed,bio}*t}+M_{stem,0}e^{v_{stem,bio}*t}$$

Generalizing the differential equation findings for the above equation gives:

$$\frac{dM_{plant}}{dt}=\sum_{tissues} \left[ M_{tissue,0}*e^{v_{tissue,bio}*t}*\left( v_{tissue,bio}+t*\frac{d}{dt}\left( v_{tissue,bio} \right) \right) \right]$$

This might be simplified by acknowledging that the time step should be equal to one, so:

$$\frac{dM_{plant}}{dt}=\sum_{tissues} \left[ M_{tissue,0}*e^{v_{tissue,bio}}*\left( v_{tissue,bio}+\frac{d}{dt}\left( v_{tissue,bio} \right) \right) \right]$$

Expanding this yields:

$$\frac{dM_{plant}}{dt}=M_{leaf,0}*e^{v_{leaf,bio}}*\left( v_{leaf,bio}+\frac{d}{dt}\left( v_{leaf,bio} \right) \right)+M_{root,0}*e^{v_{root,bio}}*\left( v_{root,bio}+\frac{d}{dt}\left( v_{root,bio} \right) \right)+M_{seed,0}*e^{v_{seed,bio}}*\left( v_{seed,bio}+\frac{d}{dt}\left( v_{seed,bio} \right) \right)+M_{stem,0}*e^{v_{stem,bio}}*\left( v_{stem,bio}+\frac{d}{dt}\left( v_{stem,bio} \right) \right)$$

This can be redefined using the following relations between each tissues growth rate and that of the leaf growth rate. These are the relations:

$$v_{root,bio}=\ln\left( \frac{x_{root}M_{leaf,0}}{x_{leaf}M_{root,0}} \right)+v_{leaf,bio}$$

$$v_{seed,bio}=ln\left( \frac{x_{seed}M_{leaf,0}}{x_{leaf}M_{seed,0}} \right)+v_{leaf,bio}$$

$$v_{stem,bio}=\ln\left( \frac{x_{stem}M_{leaf},0}{x_{leaf}M_{stem,0}} \right)+v_{leaf,bio}$$

Making these substitutions yields:

$$\frac{dM_{plant}}{dt}=M_{leaf,0}*e^{v_{leaf,bio}}*\left( v_{leaf,bio}+\frac{d}{dt}\left( v_{leaf,bio} \right) \right)+M_{root,0}*e^{\ln\left( \frac{x_{root}M_{leaf,0}}{x_{leaf}M_{root,0}} \right)+v_{leaf,bio}}*\left( \ln\left( \frac{x_{root}M_{leaf,0}}{x_{leaf}M_{root,0}} \right)+v_{leaf,bio}+\frac{d}{dt}\left( \ln\left( \frac{x_{root}M_{leaf,0}}{x_{leaf}M_{root,0}} \right)+v_{leaf,bio} \right) \right)+M_{seed,0}*e^{ln\left( \frac{x_{seed}M_{leaf,0}}{x_{leaf}M_{seed,0}} \right)+v_{leaf,bio}}*\left( ln\left( \frac{x_{seed}M_{leaf,0}}{x_{leaf}M_{seed,0}} \right)+v_{leaf,bio}+\frac{d}{dt}\left( ln\left( \frac{x_{seed}M_{leaf,0}}{x_{leaf}M_{seed,0}} \right)+v_{leaf,bio} \right) \right)+M_{stem,0}*e^{\ln\left( \frac{x_{stem}M_{leaf,0}}{x_{leaf}M_{stem,0}} \right)+v_{leaf,bio}}*\left( \ln\left( \frac{x_{stem}M_{leaf,0}}{x_{leaf}M_{stem,0}} \right)+v_{leaf,bio}+\frac{d}{dt}\left( \ln\left( \frac{x_{stem}M_{leaf,0}}{x_{leaf}M_{stem,0}} \right)+v_{leaf,bio} \right) \right)$$

Applying exponential rules and simplifying the above result gives:

$$\frac{dM_{plant}}{dt}=M_{leaf,0}*e^{v_{leaf,bio}}*\left( v_{leaf,bio}+\frac{d}{dt}\left( v_{leaf,bio} \right) \right)+M_{root,0}e^{v_{leaf,bio}}e^{\ln\left( \frac{x_{root}M_{leaf,0}}{x_{leaf}M_{root,0}} \right)}*\left( \ln\left( \frac{x_{root}M_{leaf,0}}{x_{leaf}M_{root,0}} \right)+v_{leaf,bio}+\frac{d}{dt}\left( \ln\left( \frac{x_{root}M_{leaf,0}}{x_{leaf}M_{root,0}} \right)+v_{leaf,bio} \right) \right)+M_{seed,0}*e^{v_{leaf,bio}}e^{ln\left( \frac{x_{seed}M_{leaf,0}}{x_{leaf}M_{seed,0}} \right)}*\left( ln\left( \frac{x_{seed}M_{leaf,0}}{x_{leaf}M_{seed,0}} \right)+v_{leaf,bio}+\frac{d}{dt}\left( ln\left( \frac{x_{seed}M_{leaf,0}}{x_{leaf}M_{seed,0}} \right)+v_{leaf,bio} \right) \right)+M_{stem,0}*e^{v_{leaf,bio}}e^{\ln\left( \frac{x_{stem}M_{leaf,0}}{x_{leaf}M_{stem,0}} \right)}*\left( \ln\left( \frac{x_{stem}M_{leaf,0}}{x_{leaf}M_{stem,0}} \right)+v_{leaf,bio}+\frac{d}{dt}\left( \ln\left( \frac{x_{stem}M_{leaf,0}}{x_{leaf}M_{stem,0}} \right)+v_{leaf,bio} \right) \right)$$

$$\frac{dM_{plant}}{dt}=M_{leaf,0}*e^{v_{leaf,bio}}*\left( v_{leaf,bio}+\frac{d}{dt}\left( v_{leaf,bio} \right) \right)+M_{root,0}\frac{x_{root}M_{leaf,0}}{x_{leaf}M_{root,0}}e^{v_{leaf,bio}}\left( \ln\left( \frac{x_{root}M_{leaf,0}}{x_{leaf}M_{root,0}} \right)+v_{leaf,bio}+\frac{d}{dt}\left( \ln\left( \frac{x_{root}M_{leaf,0}}{x_{leaf}M_{root,0}} \right)+v_{leaf,bio} \right) \right)+M_{seed,0}\frac{x_{seed}M_{leaf,0}}{x_{leaf}M_{seed,0}}*e^{v_{leaf,bio}}\left( ln\left( \frac{x_{seed}M_{leaf,0}}{x_{leaf}M_{seed,0}} \right)+v_{leaf,bio}+\frac{d}{dt}\left( ln\left( \frac{x_{seed}M_{leaf,0}}{x_{leaf}M_{seed,0}} \right)+v_{leaf,bio} \right) \right)+M_{stem,0}\frac{x_{stem}M_{leaf,0}}{x_{leaf}M_{stem,0}}*e^{v_{leaf,bio}}\left( \ln\left( \frac{x_{stem}M_{leaf,0}}{x_{leaf}M_{stem,0}} \right)+v_{leaf,bio}+\frac{d}{dt}\left( \ln\left( \frac{x_{stem}M_{leaf,0}}{x_{leaf}M_{stem,0}} \right)+v_{leaf,bio} \right) \right)$$

$$\frac{dM_{plant}}{dt}=e^{v_{leaf,bio}}\left[ M_{leaf,0}\left( v_{leaf,bio}+\frac{d}{dt}\left( v_{leaf,bio} \right) \right)+\frac{x_{root}M_{leaf,0}}{x_{leaf}}\left( \ln\left( \frac{x_{root}M_{leaf,0}}{x_{leaf}M_{root,0}} \right)+v_{leaf,bio}+\frac{d}{dt}\left( \ln\left( \frac{x_{root}M_{leaf,0}}{x_{leaf}M_{root,0}} \right)+v_{leaf,bio} \right) \right)+\frac{x_{seed}M_{leaf,0}}{x_{leaf}}\left( ln\left( \frac{x_{seed}M_{leaf,0}}{x_{leaf}M_{seed,0}} \right)+v_{leaf,bio}+\frac{d}{dt}\left( ln\left( \frac{x_{seed}M_{leaf,0}}{x_{leaf}M_{seed,0}} \right)+v_{leaf,bio} \right) \right)+\frac{x_{stem}M_{leaf,0}}{x_{leaf}}\left( \ln\left( \frac{x_{stem}M_{leaf,0}}{x_{leaf}M_{stem,0}} \right)+v_{leaf,bio}+\frac{d}{dt}\left( \ln\left( \frac{x_{stem}M_{leaf,0}}{x_{leaf}M_{stem,0}} \right)+v_{leaf,bio} \right) \right) \right]$$

Here it might be best to define some parameters used above. The term $x_{T}$ represents the mass ratio of of tissues $T$ to overall plant mass for the next time step, in other words:

$$x_{leaf}=\frac{M_{leaf}}{M_{plant}}$$

$$x_{root}=\frac{M_{root}}{M_{plant}}$$

$$x_{seed}=\frac{M_{seed}}{M_{plant}}$$

$$x_{stem}=\frac{M_{stem}}{M_{plant}}$$

Continuing:

$$\frac{dM_{plant}}{dt}=e^{v_{leaf,bio}}M_{leaf,0}\left[ \left( v_{leaf,bio}+\frac{d}{dt}\left( v_{leaf,bio} \right) \right)+\frac{x_{root}}{x_{leaf}}\left( \ln\left( \frac{x_{root}M_{leaf,0}}{x_{leaf}M_{root,0}} \right)+v_{leaf,bio}+\frac{d}{dt}\left( \ln\left( \frac{x_{root}M_{leaf,0}}{x_{leaf}M_{root,0}} \right)+v_{leaf,bio} \right) \right)+\frac{x_{seed}}{x_{leaf}}\left( ln\left( \frac{x_{seed}M_{leaf,0}}{x_{leaf}M_{seed,0}} \right)+v_{leaf,bio}+\frac{d}{dt}\left( ln\left( \frac{x_{seed}M_{leaf,0}}{x_{leaf}M_{seed,0}} \right)+v_{leaf,bio} \right) \right)+\frac{x_{stem}}{x_{leaf}}\left( \ln\left( \frac{x_{stem}M_{leaf,0}}{x_{leaf}M_{stem,0}} \right)+v_{leaf,bio}+\frac{d}{dt}\left( \ln\left( \frac{x_{stem}M_{leaf,0}}{x_{leaf}M_{stem,0}} \right)+v_{leaf,bio} \right) \right) \right]$$

$$\frac{dM_{plant}}{dt}=e^{v_{leaf,bio}}M_{leaf,0}\left[ \left( v_{leaf,bio}+\frac{d}{dt}\left( v_{leaf,bio} \right) \right)+\left( \frac{x_{root}}{x_{leaf}}\ln\left( \frac{x_{root}M_{leaf,0}}{x_{leaf}M_{root,0}} \right)+\frac{x_{root}}{x_{leaf}}v_{leaf,bio}+\frac{x_{root}}{x_{leaf}}\frac{d}{dt}\left( \ln\left( \frac{x_{root}M_{leaf,0}}{x_{leaf}M_{root,0}} \right)+v_{leaf,bio} \right) \right)+\left( \frac{x_{seed}}{x_{leaf}}ln\left( \frac{x_{seed}M_{leaf,0}}{x_{leaf}M_{seed,0}} \right)+\frac{x_{seed}}{x_{leaf}}v_{leaf,bio}+\frac{x_{seed}}{x_{leaf}}\frac{d}{dt}\left( ln\left( \frac{x_{seed}M_{leaf,0}}{x_{leaf}M_{seed,0}} \right)+v_{leaf,bio} \right) \right)+\left( \frac{x_{stem}}{x_{leaf}}\ln\left( \frac{x_{stem}M_{leaf,0}}{x_{leaf}M_{stem,0}} \right)+\frac{x_{stem}}{x_{leaf}}v_{leaf,bio}+\frac{x_{stem}}{x_{leaf}}\frac{d}{dt}\left( \ln\left( \frac{x_{stem}M_{leaf,0}}{x_{leaf}M_{stem,0}} \right)+v_{leaf,bio} \right) \right) \right]$$

$$\frac{dM_{plant}}{dt}=e^{v_{leaf,bio}}M_{leaf,0}\left[ v_{leaf,bio}+\frac{d}{dt}\left( v_{leaf,bio} \right)+\frac{x_{root}}{x_{leaf}}\ln\left( \frac{x_{root}M_{leaf,0}}{x_{leaf}M_{root,0}} \right)+\frac{x_{root}}{x_{leaf}}v_{leaf,bio}+\frac{x_{root}}{x_{leaf}}\frac{d}{dt}\left( \ln\left( \frac{x_{root}M_{leaf,0}}{x_{leaf}M_{root,0}} \right)+v_{leaf,bio} \right)+\frac{x_{seed}}{x_{leaf}}ln\left( \frac{x_{seed}M_{leaf,0}}{x_{leaf}M_{seed,0}} \right)+\frac{x_{seed}}{x_{leaf}}v_{leaf,bio}+\frac{x_{seed}}{x_{leaf}}\frac{d}{dt}\left( ln\left( \frac{x_{seed}M_{leaf,0}}{x_{leaf}M_{seed,0}} \right)+v_{leaf,bio} \right)+\frac{x_{stem}}{x_{leaf}}\ln\left( \frac{x_{stem}M_{leaf,0}}{x_{leaf}M_{stem,0}} \right)+\frac{x_{stem}}{x_{leaf}}v_{leaf,bio}+\frac{x_{stem}}{x_{leaf}}\frac{d}{dt}\left( \ln\left( \frac{x_{stem}M_{leaf,0}}{x_{leaf}M_{stem,0}} \right)+v_{leaf,bio} \right) \right]$$

$$\frac{dM_{plant}}{dt}=e^{v_{leaf,bio}}M_{leaf,0}\left[ v_{leaf,bio}+\frac{x_{root}}{x_{leaf}}\ln\left( \frac{x_{root}M_{leaf,0}}{x_{leaf}M_{root,0}} \right)+\frac{x_{root}}{x_{leaf}}v_{leaf,bio}+\frac{x_{seed}}{x_{leaf}}ln\left( \frac{x_{seed}M_{leaf,0}}{x_{leaf}M_{seed,0}} \right)+\frac{x_{seed}}{x_{leaf}}v_{leaf,bio}+\frac{x_{stem}}{x_{leaf}}\ln\left( \frac{x_{stem}M_{leaf,0}}{x_{leaf}M_{stem,0}} \right)+\frac{x_{stem}}{x_{leaf}}v_{leaf,bio}+\frac{d}{dt}\left( v_{leaf,bio} \right)+\frac{x_{root}}{x_{leaf}}\frac{d}{dt}\left( \ln\left( \frac{x_{root}M_{leaf,0}}{x_{leaf}M_{root,0}} \right)+v_{leaf,bio} \right)+\frac{x_{seed}}{x_{leaf}}\frac{d}{dt}\left( ln\left( \frac{x_{seed}M_{leaf,0}}{x_{leaf}M_{seed,0}} \right)+v_{leaf,bio} \right)+\frac{x_{stem}}{x_{leaf}}\frac{d}{dt}\left( \ln\left( \frac{x_{stem}M_{leaf,0}}{x_{leaf}M_{stem,0}} \right)+v_{leaf,bio} \right) \right]$$

Combine derivative terms for the sake of combining like terms:

$$\frac{dM_{plant}}{dt}=e^{v_{leaf,bio}}M_{leaf,0}\left[ v_{leaf,bio}+\frac{x_{root}}{x_{leaf}}\ln\left( \frac{x_{root}M_{leaf,0}}{x_{leaf}M_{root,0}} \right)+\frac{x_{root}}{x_{leaf}}v_{leaf,bio}+\frac{x_{seed}}{x_{leaf}}ln\left( \frac{x_{seed}M_{leaf,0}}{x_{leaf}M_{seed,0}} \right)+\frac{x_{seed}}{x_{leaf}}v_{leaf,bio}+\frac{x_{stem}}{x_{leaf}}\ln\left( \frac{x_{stem}M_{leaf,0}}{x_{leaf}M_{stem,0}} \right)+\frac{x_{stem}}{x_{leaf}}v_{leaf,bio}+\frac{d}{dt}\left( v_{leaf,bio}+\frac{x_{root}}{x_{leaf}}\ln\left( \frac{x_{root}M_{leaf,0}}{x_{leaf}M_{root,0}} \right)+\frac{x_{root}}{x_{leaf}}v_{leaf,bio}+\frac{x_{seed}}{x_{leaf}}ln\left( \frac{x_{seed}M_{leaf,0}}{x_{leaf}M_{seed,0}} \right)+\frac{x_{seed}}{x_{leaf}}v_{leaf,bio}+\frac{x_{stem}}{x_{leaf}}\ln\left( \frac{x_{stem}M_{leaf,0}}{x_{leaf}M_{stem,0}} \right)+\frac{x_{stem}}{x_{leaf}}v_{leaf,bio} \right) \right]$$

For the sake of compactness, let us make the following substitutions:

$$\mu(t)=\frac{x_{root}M_{leaf,0}}{x_{leaf}M_{root,0}}$$

$$\theta(t)=\frac{x_{seed}M_{leaf,0}}{x_{leaf}M_{seed,0}}$$

$$\lambda(t)=\frac{x_{stem}M_{leaf,0}}{x_{leaf}M_{stem,0}}$$

Making the above equation:

$$\frac{dM_{plant}}{dt}=e^{v_{leaf,bio}}M_{leaf,0}\left[ v_{leaf,bio}+\frac{x_{root}}{x_{leaf}}\ln\left( \mu(t) \right)+\frac{x_{root}}{x_{leaf}}v_{leaf,bio}+\frac{x_{seed}}{x_{leaf}}ln\left( \theta(t) \right)+\frac{x_{seed}}{x_{leaf}}v_{leaf,bio}+\frac{x_{stem}}{x_{leaf}}\ln\left( \lambda(t) \right)+\frac{x_{stem}}{x_{leaf}}v_{leaf,bio}+\frac{d}{dt}\left( v_{leaf,bio}+\frac{x_{root}}{x_{leaf}}\ln\left( \mu(t) \right)+\frac{x_{root}}{x_{leaf}}v_{leaf,bio}+\frac{x_{seed}}{x_{leaf}}ln\left( \theta(t) \right)+\frac{x_{seed}}{x_{leaf}}v_{leaf,bio}+\frac{x_{stem}}{x_{leaf}}\ln\left( \lambda(t) \right)+\frac{x_{stem}}{x_{leaf}}v_{leaf,bio} \right) \right]$$

For the sake of clarity, let us consider only the derivative term:

$$\frac{d}{dt}\left( v_{leaf,bio}+\frac{x_{root}}{x_{leaf}}\ln\left( \mu(t) \right)+\frac{x_{root}}{x_{leaf}}v_{leaf,bio}+\frac{x_{seed}}{x_{leaf}}ln\left( \theta(t) \right)+\frac{x_{seed}}{x_{leaf}}v_{leaf,bio}+\frac{x_{stem}}{x_{leaf}}\ln\left( \lambda(t) \right)+\frac{x_{stem}}{x_{leaf}}v_{leaf,bio} \right)$$

Distributing the derivative:

$$\frac{d}{dt}\left( v_{leaf,bio} \right)+\frac{d}{dt}\left( \frac{x_{root}}{x_{leaf}}\ln\left( \mu(t) \right) \right)+\frac{d}{dt}\left( \frac{x_{root}}{x_{leaf}}v_{leaf,bio} \right)+\frac{d}{dt}\left( \frac{x_{seed}}{x_{leaf}}ln\left( \theta(t) \right) \right)+\frac{d}{dt}\left( \frac{x_{seed}}{x_{leaf}}v_{leaf,bio} \right)+\frac{d}{dt}\left( \frac{x_{stem}}{x_{leaf}}\ln\left( \lambda(t) \right) \right)+\frac{d}{dt}\left( \frac{x_{stem}}{x_{leaf}}v_{leaf,bio} \right)$$

Using the product rule:

$$\frac{d}{dt}\left( v_{leaf,bio} \right)+\frac{x_{root}}{x_{leaf}}\frac{d}{dt}\left( \ln\left( \mu(t) \right) \right)+\ln\left( \mu(t) \right)\frac{d}{dt}\left( \frac{x_{root}}{x_{leaf}} \right)+\frac{x_{root}}{x_{leaf}}\frac{d}{dt}\left( v_{leaf,bio} \right)+v_{leaf,bio}\frac{d}{dt}\left( \frac{x_{root}}{x_{leaf}} \right)+\frac{x_{seed}}{x_{leaf}}\frac{d}{dt}\left( ln\left( \theta(t) \right) \right)+ln\left( \theta(t) \right)\frac{d}{dt}\left( \frac{x_{seed}}{x_{leaf}} \right)+\frac{x_{seed}}{x_{leaf}}\frac{d}{dt}\left( v_{leaf,bio} \right)+v_{leaf,bio}\frac{d}{dt}\left( \frac{x_{seed}}{x_{leaf}} \right)+\frac{x_{stem}}{x_{leaf}}\frac{d}{dt}\left( \ln\left( \lambda(t) \right) \right)+\ln\left( \lambda(t) \right)\frac{d}{dt}\left( \frac{x_{stem}}{x_{leaf}} \right)+\frac{x_{stem}}{x_{leaf}}\frac{d}{dt}\left( v_{leaf,bio} \right)+v_{leaf,bio}\frac{d}{dt}\left( \frac{x_{stem}}{x_{leaf}} \right)$$

Combining like terms:

$$\frac{x_{leaf}}{x_{leaf}}\frac{d}{dt}\left( v_{leaf,bio} \right)+\frac{x_{root}}{x_{leaf}}\frac{d}{dt}\left( v_{leaf,bio} \right)+\frac{x_{seed}}{x_{leaf}}\frac{d}{dt}\left( v_{leaf,bio} \right)+\frac{x_{stem}}{x_{leaf}}\frac{d}{dt}\left( v_{leaf,bio} \right)+\ln\left( \mu(t) \right)\frac{d}{dt}\left( \frac{x_{root}}{x_{leaf}} \right)+v_{leaf,bio}\frac{d}{dt}\left( \frac{x_{root}}{x_{leaf}} \right)+ln\left( \theta(t) \right)\frac{d}{dt}\left( \frac{x_{seed}}{x_{leaf}} \right)+v_{leaf,bio}\frac{d}{dt}\left( \frac{x_{seed}}{x_{leaf}} \right)+\ln\left( \lambda(t) \right)\frac{d}{dt}\left( \frac{x_{stem}}{x_{leaf}} \right)+v_{leaf,bio}\frac{d}{dt}\left( \frac{x_{stem}}{x_{leaf}} \right)+\frac{x_{root}}{x_{leaf}}\frac{d}{dt}\left( \ln\left( \mu(t) \right) \right)+\frac{x_{seed}}{x_{leaf}}\frac{d}{dt}\left( ln\left( \theta(t) \right) \right)+\frac{x_{stem}}{x_{leaf}}\frac{d}{dt}\left( \ln\left( \lambda(t) \right) \right)$$

$$\left( \frac{x_{leaf}}{x_{leaf}}+\frac{x_{root}}{x_{leaf}}+\frac{x_{seed}}{x_{leaf}}+\frac{x_{stem}}{x_{leaf}} \right)\frac{d}{dt}\left( v_{leaf,bio} \right)+\left( \ln\left( \mu(t) \right)+v_{leaf,bio} \right)\frac{d}{dt}\left( \frac{x_{root}}{x_{leaf}} \right)+l\left( n\left( \theta(t) \right)+v_{leaf,bio} \right)\frac{d}{dt}\left( \frac{x_{seed}}{x_{leaf}} \right)+\left( \ln\left( \lambda(t) \right)+v_{leaf,bio} \right)\frac{d}{dt}\left( \frac{x_{stem}}{x_{leaf}} \right)+\frac{1}{x_{leaf}}\left( x_{root}\frac{d}{dt}\left( \ln\left( \mu(t) \right) \right)+x_{seed}\frac{d}{dt}\left( ln\left( \theta(t) \right) \right)+x_{stem}\frac{d}{dt}\left( \ln\left( \lambda(t) \right) \right) \right)$$

$$\frac{1}{x_{leaf}}\left( x_{leaf}+x_{root}+x_{seed}+x_{stem} \right)\frac{d}{dt}\left( v_{leaf,bio} \right)+\left( \ln\left( \mu(t) \right)+v_{leaf,bio} \right)\frac{d}{dt}\left( \frac{x_{root}}{x_{leaf}} \right)+\left( ln\left( \theta(t) \right)+v_{leaf,bio} \right)\frac{d}{dt}\left( \frac{x_{seed}}{x_{leaf}} \right)+\left( \ln\left( \lambda(t) \right)+v_{leaf,bio} \right)\frac{d}{dt}\left( \frac{x_{stem}}{x_{leaf}} \right)+\frac{1}{x_{leaf}}\left( x_{root}\frac{d}{dt}\left( \ln\left( \mu(t) \right) \right)+x_{seed}\frac{d}{dt}\left( ln\left( \theta(t) \right) \right)+x_{stem}\frac{d}{dt}\left( \ln\left( \lambda(t) \right) \right) \right)$$

Since there are only four tissues and $x_{T}$ is the mass fraction of that tissue in the total plant:

$$x_{leaf}+x_{root}+x_{seed}+x_{stem}=1$$

Therefore:

$$\frac{1}{x_{leaf}}\frac{d}{dt}\left( v_{leaf,bio} \right)+\left( \ln\left( \mu(t) \right)+v_{leaf,bio} \right)\frac{d}{dt}\left( \frac{x_{root}}{x_{leaf}} \right)+\left( ln\left( \theta(t) \right)+v_{leaf,bio} \right)\frac{d}{dt}\left( \frac{x_{seed}}{x_{leaf}} \right)+\left( \ln\left( \lambda(t) \right)+v_{leaf,bio} \right)\frac{d}{dt}\left( \frac{x_{stem}}{x_{leaf}} \right)+\frac{1}{x_{leaf}}\left( x_{root}\frac{d}{dt}\left( \ln\left( \mu(t) \right) \right)+x_{seed}\frac{d}{dt}\left( ln\left( \theta(t) \right) \right)+x_{stem}\frac{d}{dt}\left( \ln\left( \lambda(t) \right) \right) \right)$$

Further, each tissues mass fraction is a linear function of some overarching variable $s$, representing the fraction of seeds present in the plant mass compared to the maximum fraction of seed mass. This variable is generally called “seeding”. These relations are:

$$x_{leaf}=c_{leaf}s+x_{leaf,0}$$

$$x_{root}=c_{root}s+x_{root,0}$$

$$x_{seed}=c_{seed}s$$

$$x_{stem}=c_{stem}s+x_{stem,0}$$

Making the appropriate substitutions:

$$\frac{1}{x_{leaf}}\frac{d}{dt}\left( v_{leaf,bio} \right)+\left( \ln\left( \mu(t) \right)+v_{leaf,bio} \right)\frac{d}{dt}\left( \frac{c_{root}s+x_{root,0}}{c_{leaf}s+x_{leaf,0}} \right)+\left( ln\left( \theta(t) \right)+v_{leaf,bio} \right)\frac{d}{dt}\left( \frac{c_{seed}s}{c_{leaf}s+x_{leaf,0}} \right)+\left( \ln\left( \lambda(t) \right)+v_{leaf,bio} \right)\frac{d}{dt}\left( \frac{c_{stem}s+x_{stem,0}}{c_{leaf}s+x_{leaf,0}} \right)+\frac{1}{x_{leaf}}\left( x_{root}\frac{d}{dt}\left( \ln\left( \mu(t) \right) \right)+x_{seed}\frac{d}{dt}\left( ln\left( \theta(t) \right) \right)+x_{stem}\frac{d}{dt}\left( \ln\left( \lambda(t) \right) \right) \right)$$

For now, consider only the derivative terms with variable $s$ in them to avoid excessive repetitive writing. Let us look at the first such term:

$$\frac{d}{dt}\left( \frac{c_{root}s+x_{root,0}}{c_{leaf}s+x_{leaf,0}} \right)$$

Applying the quotient rule:

$$\frac{d}{dt}\left( \frac{c_{root}s+x_{root,0}}{c_{leaf}s+x_{leaf,0}} \right)=\frac{\left( c_{leaf}s+x_{leaf,0} \right)\frac{d}{dt}\left( c_{root}s+x_{root,0} \right)-\left( c_{root}s+x_{root,0} \right)\frac{d}{dt}\left( c_{leaf}s+x_{leaf,0} \right)}{\left( c_{leaf}s+x_{leaf,0} \right)^{2}}$$

Simplifying by recognizing $x_{root,0}$, $x_{leaf,0}$, $c_{root}$, $c_{leaf}$ are constants:

$$\frac{d}{dt}\left( \frac{c_{root}s+x_{root,0}}{c_{leaf}s+x_{leaf,0}} \right)=\frac{c_{root}\left( c_{leaf}s+x_{leaf,0} \right)\frac{d}{dt}\left( s \right)-c_{leaf}\left( c_{root}s+x_{root,0} \right)\frac{d}{dt}\left( s \right)}{c_{leaf}^{2}s^{2}+2c_{leaf}sx_{leaf,0}+x_{leaf,0}^{2}}$$

Combining like terms:

$$\frac{d}{dt}\left( \frac{c_{root}s+x_{root,0}}{c_{leaf}s+x_{leaf,0}} \right)=\frac{c_{root}\left( c_{leaf}s+x_{leaf,0} \right)-c_{leaf}\left( c_{root}s+x_{root,0} \right)}{c_{leaf}^{2}s^{2}+2c_{leaf}sx_{leaf,0}+x_{leaf,0}^{2}}\frac{d}{dt}\left( s \right)$$

For compactness, let us define:

$$\zeta=\frac{c_{root}\left( c_{leaf}s+x_{leaf,0} \right)-c_{leaf}\left( c_{root}s+x_{root,0} \right)}{c_{leaf}^{2}s^{2}+2c_{leaf}sx_{leaf,0}+x_{leaf,0}^{2}}$$

Therefore:

$$\frac{d}{dt}\left( \frac{c_{root}s+x_{root,0}}{c_{leaf}s+x_{leaf,0}} \right)=\zeta\frac{d}{dt}\left( s \right)$$

Similarly:

$$\frac{d}{dt}\left( \frac{c_{seed}s}{c_{leaf}s+x_{leaf,0}} \right)=\frac{\left( c_{leaf}s+x_{leaf,0} \right)\frac{d}{dt}\left( c_{seed}s \right)-\left( c_{seed}s \right)\frac{d}{dt}\left( c_{leaf}s+x_{leaf,0} \right)}{\left( c_{leaf}s+x_{leaf,0} \right)^{2}}$$

$$\frac{d}{dt}\left( \frac{c_{seed}s}{c_{leaf}s+x_{leaf,0}} \right)=\frac{c_{seed}\left( c_{leaf}s+x_{leaf,0} \right)\frac{d}{dt}\left( s \right)-c_{leaf}\left( c_{seed}s \right)\frac{d}{dt}\left( s \right)}{c_{leaf}^{2}s^{2}+2c_{leaf}sx_{leaf,0}+x_{leaf,0}^{2}}$$

$$\frac{d}{dt}\left( \frac{c_{seed}s}{c_{leaf}s+x_{leaf,0}} \right)=\frac{c_{seed}\left( c_{leaf}s+x_{leaf,0} \right)+c_{leaf}\left( c_{seed}s \right)}{c_{leaf}^{2}s^{2}+2c_{leaf}sx_{leaf,0}+x_{leaf,0}^{2}}\frac{d}{dt}\left( s \right)$$

Let:

$$\rho=\frac{c_{seed}\left( c_{leaf}s+x_{leaf,0} \right)-c_{leaf}\left( c_{seed}s \right)}{c_{leaf}^{2}s^{2}+2c_{leaf}sx_{leaf,0}+x_{leaf,0}^{2}}$$

Then:

$$\frac{d}{dt}\left( \frac{c_{seed}s}{c_{leaf}s+x_{leaf,0}} \right)=\rho\frac{d}{dt}\left( s \right)$$

And finally:

$$\frac{d}{dt}\left( \frac{c_{stem}s+x_{stem,0}}{c_{leaf}s+x_{leaf,0}} \right)=\frac{\left( c_{leaf}s+x_{leaf,0} \right)\frac{d}{dt}\left( c_{stem}s+x_{stem,0} \right)-\left( c_{stem}s+x_{stem,0} \right)\frac{d}{dt}\left( c_{leaf}s+x_{leaf,0} \right)}{\left( c_{leaf}s+x_{leaf,0} \right)^{2}}$$

$$\frac{d}{dt}\left( \frac{c_{stem}s+x_{stem,0}}{c_{leaf}s+x_{leaf,0}} \right)=\frac{c_{stem}\left( c_{leaf}s+x_{leaf,0} \right)\frac{d}{dt}\left( s \right)-c_{leaf}\left( c_{stem}s+x_{stem,0} \right)\frac{d}{dt}\left( s \right)}{c_{leaf}^{2}s^{2}+2c_{leaf}sx_{leaf,0}+x_{leaf,0}^{2}}$$

$$\frac{d}{dt}\left( \frac{c_{stem}s+x_{stem,0}}{c_{leaf}s+x_{leaf,0}} \right)=\frac{c_{stem}\left( c_{leaf}s+x_{leaf,0} \right)-c_{leaf}\left( c_{stem}s+x_{stem,0} \right)}{c_{leaf}^{2}s^{2}+2c_{leaf}sx_{leaf,0}+x_{leaf,0}^{2}}\frac{d}{dt}\left( s \right)$$

Let:

$$\iota=\frac{c_{stem}\left( c_{leaf}s+x_{leaf,0} \right)-c_{leaf}\left( c_{stem}s+x_{stem,0} \right)}{c_{leaf}^{2}s^{2}+2c_{leaf}sx_{leaf,0}+x_{leaf,0}^{2}}$$

Then:

$$\frac{d}{dt}\left( \frac{c_{stem}s+x_{stem,0}}{c_{leaf}s+x_{leaf,0}} \right)=\omega\frac{d}{dt}\left( s \right)$$

Making the appropriate substitutions:

$$\frac{1}{x_{leaf}}\frac{d}{dt}\left( v_{leaf,bio} \right)+\left( \ln\left( \mu(t) \right)+v_{leaf,bio} \right)\zeta\frac{d}{dt}\left( s \right)+\left( ln\left( \theta(t) \right)+v_{leaf,bio} \right)\rho\frac{d}{dt}\left( s \right)+\left( \ln\left( \lambda(t) \right)+v_{leaf,bio} \right)\omega\frac{d}{dt}\left( s \right)+\frac{1}{x_{leaf}}\left( x_{root}\frac{d}{dt}\left( \ln\left( \mu(t) \right) \right)+x_{seed}\frac{d}{dt}\left( ln\left( \theta(t) \right) \right)+x_{stem}\frac{d}{dt}\left( \ln\left( \lambda(t) \right) \right) \right)$$

Combining like terms:

$$\frac{1}{x_{leaf}}\frac{d}{dt}\left( v_{leaf,bio} \right)+\left( \left( \ln\left( \mu(t) \right)+v_{leaf,bio} \right)\zeta+\left( ln\left( \theta(t) \right)+v_{leaf,bio} \right)\rho+\left( \ln\left( \lambda(t) \right)+v_{leaf,bio} \right)\omega\right)\frac{d}{dt}\left( s \right)+\frac{1}{x_{leaf}}\left( x_{root}\frac{d}{dt}\left( \ln\left( \mu(t) \right) \right)+x_{seed}\frac{d}{dt}\left( ln\left( \theta(t) \right) \right)+x_{stem}\frac{d}{dt}\left( \ln\left( \lambda(t) \right) \right) \right)$$

Substituting the derivative term back into the overall equation gives:

$$\frac{dM_{plant}}{dt}=e^{v_{leaf,bio}}M_{leaf,0}\left[ v_{leaf,bio}+\frac{x_{root}}{x_{leaf}}\ln\left( \mu(t) \right)+\frac{x_{root}}{x_{leaf}}v_{leaf,bio}+\frac{x_{seed}}{x_{leaf}}ln\left( \theta(t) \right)+\frac{x_{seed}}{x_{leaf}}v_{leaf,bio}+\frac{x_{stem}}{x_{leaf}}\ln\left( \lambda(t) \right)+\frac{x_{stem}}{x_{leaf}}v_{leaf,bio}+\frac{1}{x_{leaf}}\frac{d}{dt}\left( v_{leaf,bio} \right)+\left( \left( \ln\left( \mu(t) \right)+v_{leaf,bio} \right)\zeta+\left( ln\left( \theta(t) \right)+v_{leaf,bio} \right)\rho+\left( \ln\left( \lambda(t) \right)+v_{leaf,bio} \right)\omega\right)\frac{d}{dt}\left( s \right)+\frac{1}{x_{leaf}}\left( x_{root}\frac{d}{dt}\left( \ln\left( \mu(t) \right) \right)+x_{seed}\frac{d}{dt}\left( ln\left( \theta(t) \right) \right)+x_{stem}\frac{d}{dt}\left( \ln\left( \lambda(t) \right) \right) \right) \right]$$

$$\frac{dM_{plant}}{dt}=\frac{e^{v_{leaf,bio}}M_{leaf,0}}{x_{leaf}}\left[ x_{leaf}v_{leaf,bio}+x_{root}\ln\left( \mu(t) \right)+x_{root}v_{leaf,bio}+x_{seed}ln\left( \theta(t) \right)+x_{seed}v_{leaf,bio}+x_{stem}\ln\left( \lambda(t) \right)+x_{stem}v_{leaf,bio}+\frac{d}{dt}\left( v_{leaf,bio} \right)+x_{leaf}\left( \left( \ln\left( \mu(t) \right)+v_{leaf,bio} \right)\zeta+\left( ln\left( \theta(t) \right)+v_{leaf,bio} \right)\rho+\left( \ln\left( \lambda(t) \right)+v_{leaf,bio} \right)\omega\right)\frac{d}{dt}\left( s \right)+\left( x_{root}\frac{d}{dt}\left( \ln\left( \mu(t) \right) \right)+x_{seed}\frac{d}{dt}\left( ln\left( \theta(t) \right) \right)+x_{stem}\frac{d}{dt}\left( \ln\left( \lambda(t) \right) \right) \right) \right]$$

$$\frac{dM_{plant}}{dt}=\frac{e^{v_{leaf,bio}}M_{leaf,0}}{x_{leaf}}\left[ x_{leaf}v_{leaf,bio}+x_{root}\left( \ln\left( \mu(t) \right)+v_{leaf,bio} \right)+x_{seed}\left( ln\left( \theta(t) \right)+v_{leaf,bio} \right)+x_{stem}\left( \ln\left( \lambda(t) \right)+v_{leaf,bio} \right)+\frac{d}{dt}\left( v_{leaf,bio} \right)+x_{leaf}\left( \left( \ln\left( \mu(t) \right)+v_{leaf,bio} \right)\zeta+\left( ln\left( \theta(t) \right)+v_{leaf,bio} \right)\rho+\left( \ln\left( \lambda(t) \right)+v_{leaf,bio} \right)\omega\right)\frac{d}{dt}\left( s \right)+\left( x_{root}\frac{d}{dt}\left( \ln\left( \mu(t) \right) \right)+x_{seed}\frac{d}{dt}\left( ln\left( \theta(t) \right) \right)+x_{stem}\frac{d}{dt}\left( \ln\left( \lambda(t) \right) \right) \right) \right]$$

Let:

$$\psi\left( t \right)=\ln\left( \mu(t) \right)+v_{leaf,bio}$$

$$\pi\left( t \right)=ln\left( \theta(t) \right)+v_{leaf,bio}$$

$$\kappa\left( t \right)=\ln\left( \lambda(t) \right)+v_{leaf,bio}$$

Making the substitutions:

$$\frac{dM_{plant}}{dt}=\frac{e^{v_{leaf,bio}}M_{leaf,0}}{x_{leaf}}\left[ x_{leaf}v_{leaf,bio}+x_{root}\psi\left( t \right)+x_{seed}\pi\left( t \right)+x_{stem}\kappa\left( t \right)+\frac{d}{dt}\left( v_{leaf,bio} \right)+x_{leaf}\left( \psi\left( t \right)\zeta\left( t \right)+\pi\left( t \right)\rho\left( t \right)+\kappa\left( t \right)\omega\left( t \right) \right)\frac{d}{dt}\left( s \right)+x_{root}\frac{d}{dt}\left( \ln\left( \mu(t) \right) \right)+x_{seed}\frac{d}{dt}\left( ln\left( \theta(t) \right) \right)+x_{stem}\frac{d}{dt}\left( \ln\left( \lambda(t) \right) \right) \right]$$

Making a final substitution separating growth rates from equations dependent on time points:

$$\frac{dM_{plant}}{dt}=\frac{e^{v_{leaf,bio}}M_{leaf,0}}{x_{leaf}}\left[ x_{leaf}v_{leaf,bio}+\frac{d}{dt}\left( v_{leaf,bio} \right)+\xi\right]$$

$$\xi(t)=x_{root}\psi\left( t \right)+x_{seed}\pi\left( t \right)+x_{stem}\kappa\left( t \right)+x_{leaf}\left( \psi\left( t \right)\zeta\left( t \right)+\pi\left( t \right)\rho\left( t \right)+\kappa\left( t \right)\omega\left( t \right) \right)\frac{d}{dt}\left( s \right)+x_{root}\frac{d}{dt}\left( \ln\left( \mu(t) \right) \right)+x_{seed}\frac{d}{dt}\left( ln\left( \theta(t) \right) \right)+x_{stem}\frac{d}{dt}\left( \ln\left( \lambda(t) \right) \right)$$

Therefore the derivative is:

$$\frac{dM_{plant}}{dt}=\frac{e^{v_{leaf,bio}}M_{leaf,0}}{x_{leaf}}\left[ x_{leaf}v_{leaf,bio}+\frac{d}{dt}\left( v_{leaf,bio} \right)+\xi\left( t \right) \right]$$

$$\xi\left( t \right)=x_{root}\psi\left( t \right)+x_{seed}\pi\left( t \right)+x_{stem}\kappa\left( t \right)+x_{leaf}\left( \psi\left( t \right)\zeta\left( t \right)+\pi\left( t \right)\rho\left( t \right)+\kappa\left( t \right)\iota\left( t \right) \right)\frac{d}{dt}\left( s \right)+x_{root}\frac{d}{dt}\left( \ln\left( \mu(t) \right) \right)+x_{seed}\frac{d}{dt}\left( ln\left( \theta(t) \right) \right)+x_{stem}\frac{d}{dt}\left( \ln\left( \lambda(t) \right) \right)$$

$$\psi\left( t \right)=\ln\left( \mu(t) \right)+v_{leaf,bio}$$

$$\pi\left( t \right)=ln\left( \theta(t) \right)+v_{leaf,bio}$$

$$\kappa\left( t \right)=\ln\left( \lambda(t) \right)+v_{leaf,bio}$$

$$\mu(t)=\frac{x_{root}M_{leaf,0}}{x_{leaf}M_{root,0}}$$

$$\theta(t)=\frac{x_{seed}M_{leaf,0}}{x_{leaf}M_{seed,0}}$$

$$\lambda(t)=\frac{x_{stem}M_{leaf,0}}{x_{leaf}M_{stem,0}}$$

$$\zeta=\frac{c_{root}\left( c_{leaf}s+x_{leaf,0} \right)-c_{leaf}\left( c_{root}s+x_{root,0} \right)}{c_{leaf}^{2}s^{2}+2c_{leaf}sx_{leaf,0}+x_{leaf,0}^{2}}$$

$$\rho=\frac{c_{seed}\left( c_{leaf}s+x_{leaf,0} \right)-c_{leaf}\left( c_{seed}s \right)}{c_{leaf}^{2}s^{2}+2c_{leaf}sx_{leaf,0}+x_{leaf,0}^{2}}$$

$$\iota=\frac{c_{stem}\left( c_{leaf}s+x_{leaf,0} \right)-c_{leaf}\left( c_{stem}s+x_{stem,0} \right)}{c_{leaf}^{2}s^{2}+2c_{leaf}sx_{leaf,0}+x_{leaf,0}^{2}}$$

$$\frac{d}{dt}\left( v_{leaf growth, n} \right)=\frac{3v_{leaf growth,n}-4v_{leaf growth, n-1/3}+v_{leaf growth,n-2/3}}{2\left[ \frac{1}{3}h \right]}$$

All equations not dependent on growth rate will be treated as parameters at that time point, whereas those that are dependent on the biomass rate will be treated as variables. Therefore:

| Parameters | Variables |
| --- | --- |
| $x_{leaf}, x_{root}, x_{seed},x_{stem}$  $\mu\left( t \right),\theta\left( t \right),\lambda(t)$  $\zeta,\rho,\omega$ | $v_{leaf,bio}$  $\frac{d}{dt}\left( v_{leaf,bio} \right)$  $\frac{dM_{plant}}{dt}$  $\psi\left( t \right),\pi\left( t \right),\kappa\left( t \right),\xi\left( t \right)$ |

In the first few iterations, Euler’s method has been used, but has been found to be unstable for this system. Next, Kutta’s third-order rule for the Runge-Kutta method has been used; however, the negative value in the Butchers tableau occasionally has resulted in large, negative derivative estimates which could not have been overcome by the other terms. This would result in negative plant mass at the next value of $n$, which in this context was nonsense. Therefore, Huenn’s rule for the third order Runge-Kutta method has been used as it does not contain a negative value in its Butcher tableau, but still has resulted in evenly spaced steps, which has been important for equations (50), (51), and (52). Heunn’s rule for third-order Runge-Kutta is as follows:

$$k_{1}=f(\vec{x}_{n},\vec{y}_{n})$$

$$k_{2}=f\left( \vec{x}_{n}+\frac{1}{3}h,y_{n}+\frac{1}{3}hk_{1} \right)$$

$$k_{3}=f\left( \vec{x}_{n}+\frac{2}{3}h,y_{n}+\frac{2}{3}hk_{2} \right)$$

$$y_{n+1}=y_{n}+h\left( \frac{1}{4}k_{1}+\frac{3}{4}k_{3} \right)$$

Where in this model $k_{1}$ is the estimate in change in plant mass with respect to time from $n$ to $n+1/3$, $k_{2}$ is the estimate in change in plant mass with respect to time from $n+1/3$ to $n+2/3$, $k_{1}$ is the estimate in change in plant mass with respect to time from $n+2/3$ to $n+1$.

Overall plant mass has been chosen to be updated, rather than each individual tissue. This was done to avoid either solving for the change in mass with respect to time for each tissue (effectively quadrupling solve time); having the tissue ratios gradually drift due to the large number of solved points and rounding error; or having a variety of issues with the seed tissue. Specifically, the seed tissue is problematic as it transits from zero to non-zero mass and from non-zero to zero mass, but the biomass growth rates are exponential growth rates. When transitioning from zero to non-zero mass, some non-zero amount of mass would have to have been postulated as M_seed,n_ in order for a non-zero M_seed,n+1_ to result. Essentially, it requires the mass balance be violated by creating a non-zero amount of mass inside the system. It has also been difficult to determine what the assumed $M_{seed,n}$ value should be used. Similarly, it is not possible to result in M_seed,n+1_=0. Therefore, mass balance must have also been violated to remove a set amount of mass. The method used has been chosen to avoid these issues.
