## Supplemental File 23 for "A Computational Framework to Study the Primary Lifecycle Metabolism of *Arabidopsis thaliana*"

**General File Structure Used in Workflow:**

**Main Folder:** p-ath780

**Files:** ModelAnalysis.xlsx, makeGrowthInputs.pl, growthSpecsNames.txt, growthSpecs.txt, timepointsH.txt, timepoints.txt, timeData.txt, sunrise.txt, sunset.txt, timeofday.txt, p-ath780.gms, and Supplemental_File_10.docx

**Sub-Folder:** p-athLeaf

**Files:** p-ath780Leaf.txt, convert.py, pathList.csv, PathGetRxnsComps.pl, ModelPathComp.pl, RxnstoGenes.pl

**Workflow:**

1) Add all above files to folder
2) Run “convert.py” to generate files readable by and necessary for p-ath780.gms
3) If desired, run PathGetRxnsComps.pl and ModelPathComp.pl to get initial set of data behind the construction of Fig. 2.
4) Run RxnstoGenes.pl if desired to get initial GPR link information which is also used in Fig. 2.

**Sub-Folder:** p-athRoot

**Files:** p-ath780Root.txt, convert.py, pathList.csv, PathGetRxnsComps.pl, ModelPathComp.pl, RxnstoGenes.pl

**Workflow:**

1) Add all above files to folder
2) Run “convert.py” to generate files readable by and necessary for p-ath780.gms
3) If desired, run PathGetRxnsComps.pl and ModelPathComp.pl to get initial set of data behind the construction of Fig. 2.
4) Run RxnstoGenes.pl if desired to get initial GPR link information which is also used in Fig. 2.

**Sub-Folder:** p-athSeed

**Files:** p-ath780Seed.txt, convert.py, pathList.csv, PathGetRxnsComps.pl, ModelPathComp.pl, RxnstoGenes.pl

**Workflow:**

1) Add all above files to folder
2) Run “convert.py” to generate files readable by and necessary for p-ath780.gms
3) If desired, run PathGetRxnsComps.pl and ModelPathComp.pl to get initial set of data behind the construction of Fig. 2.
4) Run RxnstoGenes.pl if desired to get initial GPR link information which is also used in Fig. 2.

**Sub-Folder:** p-athStem

**Files:** p-ath780Stem.txt, convert.py, pathList.csv, PathGetRxnsComps.pl, ModelPathComp.pl, RxnstoGenes.pl

**Workflow:**

1) Add all above files to folder
2) Run “convert.py” to generate files readable by and necessary for p-ath780.gms
3) If desired, run PathGetRxnsComps.pl and ModelPathComp.pl to get initial set of data behind the construction of Fig. 2.
4) Run RxnstoGenes.pl if desired to get initial GPR link information which is also used in Fig. 2.
